# Cannabidiol rescues age-associated cognitive decline in mouse model

**DOI:** 10.64898/2026.04.12.711942

**Authors:** Anuradha Kesharwani, Devidas Lahamge, Alok Sharma, Murali Kumarasamy, Velayutham Ravichandiran, Vipan Kumar Parihar

**Author notes:** Florida Atlantic University, Stiles-Nicholson Brain Institute, Jupiter, Florida - 33458, USA. Central University of Haryana, Mahendergarh, Haryana-123031, India (Vipan Kumar Parihar). **Corresponding Author** Dr. Vipan K. Parihar, Department of Pharmacology and Toxicology, Department of Regulatory Toxicology, National Institute of Pharmaceutical Education and Research, Hajipur - 844102, Bihar, India.

## Abstract

This study addresses key gaps in our understanding of the cognitive changes associated with normal brain aging and their underlying structural and functional correlates. Declining levels of brain endocannabinoids (eCB), particularly 2-arachidonoylglycerol (2-AG), are thought to contribute to age-related cognitive impairment, but the mechanisms involved remain poorly understood. Here, we show that levels of 2-AG and its synthesizing enzyme, diacylglycerol lipase-α (DAGL-α), are significantly reduced in the medial prefrontal cortex (mPFC) of aged mice. This decline is associated with impairments in mood and memory, as demonstrated by behavioral analyses. Immunofluorescence studies further revealed reduced expression of cannabinoid receptors CB1 and CB2 on microglia in aged brains, suggesting that diminished eCB signaling may contribute to enhanced neuroinflammation. Consistent with this idea, microglia from aged mice exhibited increased HMGB1–TLR4–NF-κB signaling, indicative of a pro-inflammatory state. LC-HR-MS analysis also showed elevated glutamate levels and reduced glutamine and GABA levels in the mPFC, linking impaired eCB signaling to excitotoxic imbalance during aging. Importantly, intraperitoneal administration of cannabidiol (CBD) to aged mice reversed mood and memory deficits, restored CB1 and CB2 receptor expression, attenuated HMGB1–TLR4–NF-κB signaling, and normalized the glutamate–glutamine/GABA balance in the mPFC. Collectively, these findings identify eCB signaling as a critical regulator of age-associated cognitive decline and support CBD as a potential therapeutic strategy to promote healthy brain aging.

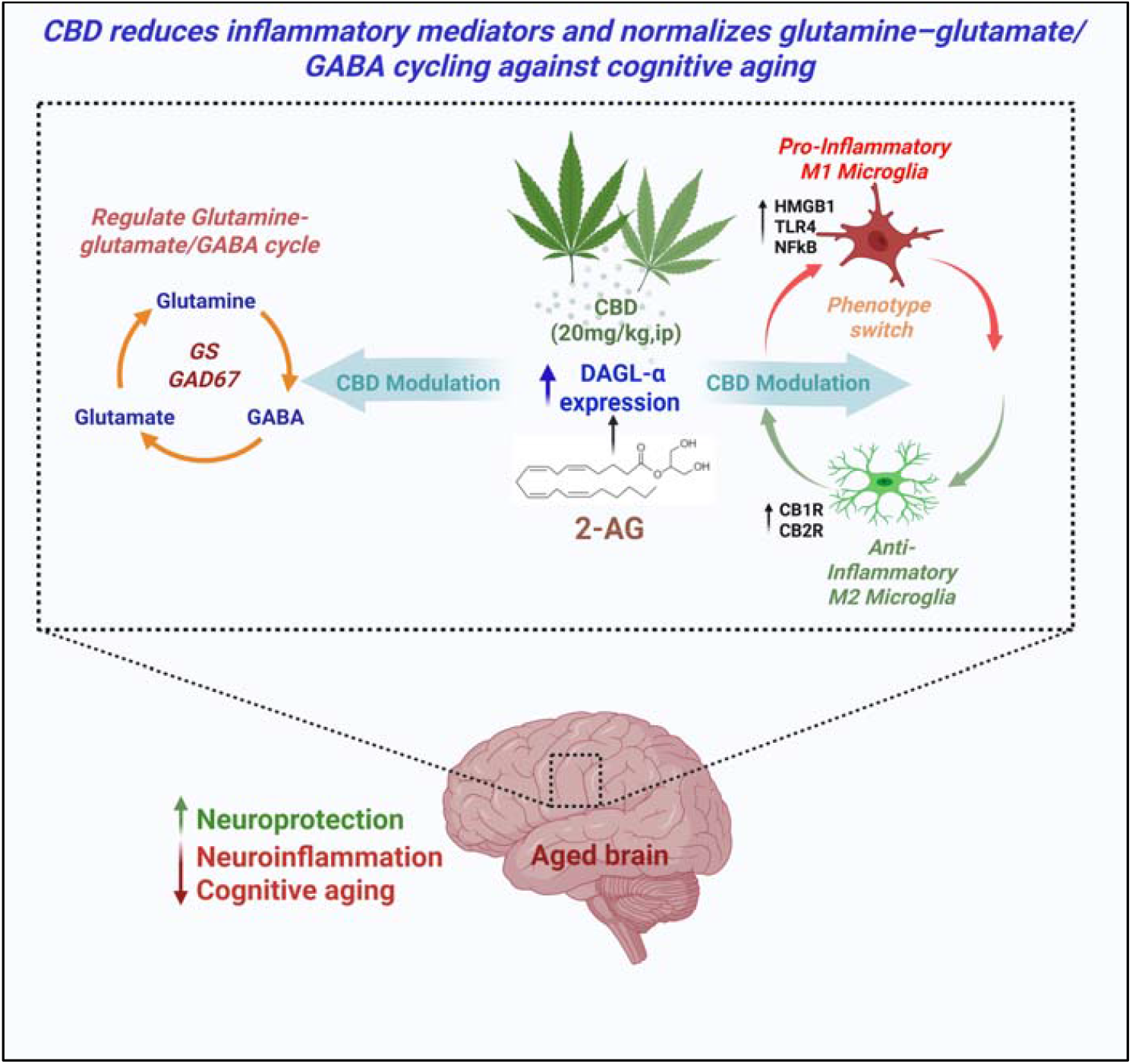

**Highlights:**

- Reduction of 2-AG in the brain’s medial prefrontal cortex (mPFC) over time is a hallmark of cognitive decline.
- Pro-inflammatory signalling such as HMGB1 (High Mobility Group Box 1)-TLR4 (Toll-like receptor 4)-NF-kB (Nuclear Factor kappa B) signalling are elevated in aged brain.
- Altered glutamine-glutamate/GABA cycle associated with reduced 2-AG levels.
- CBD administration mitigates cognitive decline associated with aging and diminishes neuroinflammation.

## Introduction

Aging is associated with a progressive decline in executive function, memory, and learning capacity, significantly impairing cognitive performance and quality of life(Idowu and Szameitat, 2023). At the biological level, normal aging involves the gradual accumulation of molecular and cellular damage, including genomic instability, cellular senescence, epigenetic alterations, impaired proteostasis, mitochondrial dysfunction, and chronic inflammation(Li et al., 2024; Panchal, 2024; Tenchov et al., 2024; Wang et al., 2022). In age-related disorders such as Alzheimer disease (AD), these processes are further exacerbated, leading to accelerated cognitive decline and profound deterioration in quality of life(Weaver, 2024). Therefore, elucidating the mechanisms underlying brain aging is essential for developing effective interventions to mitigate cognitive decline and promote healthy aging.

The endocannabinoid (eCB) signalling system is a finely tuned neuromodulatory network that maintains brain homeostasis by integrating synaptic, neuroimmune, and metabolic processes. It comprises endogenous lipid mediators, primarily 2-arachidonoylglycerol (2-AG) and anandamide (AEA), along with their biosynthetic and degradative enzymes and cannabinoid receptors, including CB1 and CB2(Bilkei-Gorzo, 2012; Di Marzo, 2015). These receptors coordinate signalling to regulate glial activity, neuronal network, and immune tone in the brain(Stella, 2009). Recent studies suggest that the eCB system could contribute to brain aging and cognitive function. For instance, mice depleted with CB1 or CB2 receptors exhibit dysregulation of neurotransmitter systems, impaired immune homeostasis, and enhanced neuroinflammation, ultimately leading to severe neurotoxicity and cognitive deficits(Springs et al., 2008; Spyridakos et al., 2022).

Despite extensive research, the role of the eCB system in aging remains incompletely understood. Here, we show that a decline in 2-AG signalling is associated with age-related cognitive impairment, in part by promoting neuroinflammation and disrupting neurotransmitter imbalance. Furthermore, we demonstrate that these age-associated changes can be reversed by cannabidiol (CBD) administration, supporting the eCB system as a viable target to improve cognitive function during aging.

## Materials and Methods

### Animals

All animal procedures conducted in this study adhered strictly to the guidelines of the Committee for the Control and Supervision of Experiments on Animals (CCSEA), Ministry of Environment and Forests, Government of India, New Delhi. The experimental protocol was reviewed and approved by the Institutional Animal Ethics Committee (IAEC) of the National Institute of Pharmaceutical Education and Research (NIPER), Hajipur, India (Approval No: NIPER-H/IAEC/21/22). Male C57BL/6J mice of different ages were procured from the ICMR-National Institute of Nutrition (NIN), Hyderabad, India. Animals were maintained under standard laboratory conditions at the Central Animal Research Facility, NIPER Hajipur, with free (ad libitum) access to food and water, with a controlled environment of 20 ± 1°C temperature, 50 ± 10% relative humidity, and a 12-hour light/dark cycle.

### Drug

Cannabidiol (CBD; 20 mg/kg, intraperitoneally; *Cayman Chemical*) was freshly prepared in 0.1% ethyl alcohol (*SRL*) immediately prior to administration to ensure stability and optimal bioavailability.

### Experimental design

To investigate age-associated alterations in eCB signalling, two separate sets of animals were utilized. The first set was employed to assess the extent and temporal progression of altered eCB signalling in relation to neurocognitive dysfunction. Mice were randomly assigned to three groups (n = 10 per group): 4-Month, 12-Month, and 20-Month aged mice were chosen to represent young adult, middle-aged, and aged mice, respectively, based on established classifications of mouse aging that correspond to different stages of cognitive and neurobiological decline. Animals from each age group were subjected to a series of behavioral tests and biochemical assessments. The behavioral assays included the Temporal Order (TO) test, Novel Object Recognition (NOR), Object-in-Place (OiP) test, and Forced Swim Test (FST) ***(Fig. 1A)***.

**Fig. 1:**
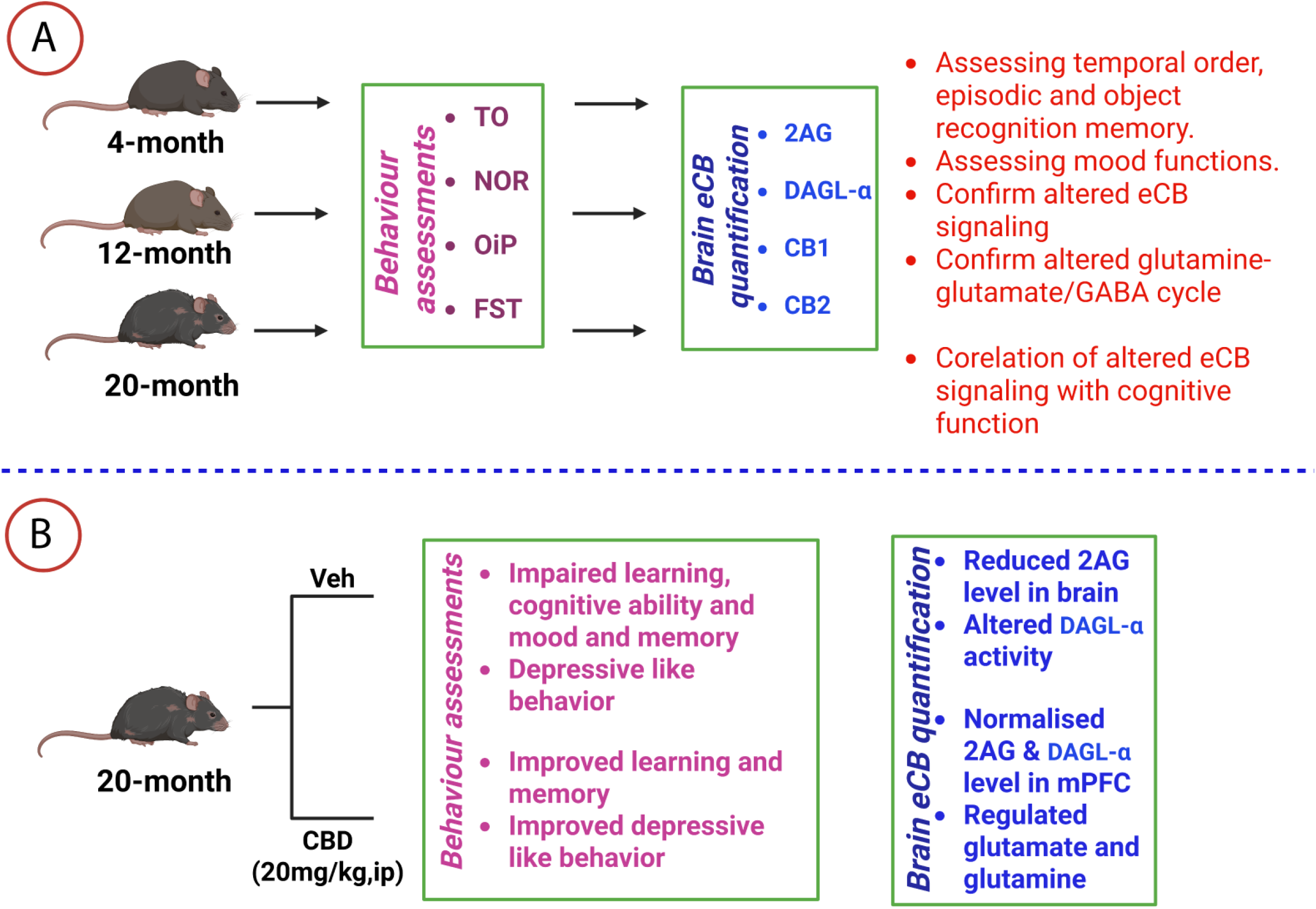
The graphic illustrates the progression of multiple experiments and analyses conducted in this study. The experimental protocol comprised two distinct sets of mice. The first set included three separate age groups: four, twelve, and twenty months old mice **(A)**. The second set consisted of 20 months + Veh and 20 months + CBD. Mice were euthanized 24 hours after the last behavioral assessment. Their brains were euthanized 24 hours after the last behavioral assessment. Their brains wereextracted from the skull and stored at –80°C for subsequent biochemical examination.

The second set of animals was used to evaluate the efficacy of CBD in mitigating eCB signalling deficits in aged mice. This set comprised two groups (n = 10 per group): 20-month-old with vehicle (20-Month + Veh) and 20-month-old with CBD treatment (20-Month + CBD). Mice in the CBD-treated group received daily intraperitoneal (ip) injections of CBD (20 mg/kg, freshly prepared in 0.1% ethyl alcohol) for a duration of two weeks, based on previously published work from our lab, as well as prior preclinical studies demonstrating its efficacy in modulating neuroinflammatory and behavioral outcomes without inducing toxicity or motor side effects(Kaplan et al., 2017). Following the treatment period, animals were evaluated for mood and memory functions using the same behavioral tests as the first set (TO, NOR, OiP, and FST; ***Fig. 1B***). All behavioral assessments were conducted during the light phase, between 09:00 and 14:00 h, to minimize circadian variability. Detailed protocols for each behavioral test are provided in ***Supplementary File 1***.

### Behavior assessment

TO test evaluates an animal’s ability to distinguish between previously explored objects based on their relative recency, reflecting the capacity to recall the temporal sequence of past experiences. This task relies on interactions between the perirhinal cortex and the medial prefrontal cortex (mPFC). The NOR test assesses an animal’s preference for novelty, which relies on the proper functioning of the mPFC. This task evaluates recognition memory by determining whether the animal spends more time exploring a novel object compared to a familiar one, indicating the ability to discriminate. The OiP task evaluates associative recognition memory, which involves the integration of information from the hippocampus, perirhinal cortex, and mPFC (Kesharwani et al., 2025a; Kesharwani et al., 2025b; Parihar et al., 2020; Parihar et al., 2021). This test examines the animal’s ability to detect changes in the spatial arrangement of familiar objects, thereby assessing its capacity to recognize and associate objects with specific locations. All objects used in this study were designed to have identical inherent preferences and were easily distinguishable by control mice. The objects were standardized in height and volume but varied in shape, texture, and appearance. To minimize potential biases related to object characteristics, the objects were randomized during the behavioral assessments. Data collection and analysis were performed by two independent observers who were blinded to the group identities of the animals. A discrimination index (DI) was calculated to evaluate the preference or indifference toward novelty by using formula given below:

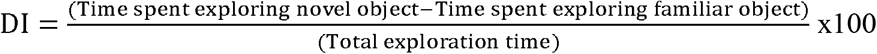

A positive DI score indicates a tendency to prefer novelty, whereas a negative score suggests a lack of this preference. Comprehensive details of the behavioral testing procedures are available in ***Supplementary File 1***.

### Tissue sample processing for subsequent biochemical and immunofluorescence (IF) evaluation

After the last behavioral test, mice were randomly divided in two different group for saline and 4% paraformaldehyde (PFA) perfusion. Mice were euthanized under deep anaesthesia using isoflurane in a plexiglas chamber, and their brains were isolated for biochemical and IF studies (Kesharwani et al., 2025a; Kesharwani et al., 2025b; Singh et al., 2025). For ELISA assays and neurotransmitter estimations, the animals were perfused with cold heparinized saline, and the brains were rapidly removed from the skull. The mPFC tissue was used for the ELISA assay, and neurotransmitter estimation. For IF, mice were perfused with 4% PFA followed by cold heparinized saline. ***Supplementary File 1*** provides a comprehensive overview of the methodologies used for ELISA assay, neurotransmitters estimation and IF analysis.

### Measurement of 2-Arachidonoylglycerol (2-AG) levels and Diacylglycerol Lipase-α (DAGL-α) enzymatic activity

The concentration of the 2-AG (Krishgen Biosystems, KLU0041) and the enzymatic activity of its synthesizing enzyme, DAGL-α (ELK Biotechnology, ELK9005), were quantified in mPFC tissue homogenates following the respective manufacturers’ protocols(Kesharwani et al., 2025a). Comprehensive experimental procedures are provided in ***Supplementary File 1***.

### Visualization via immunofluorescence (IF) and confocal imaging techniques

After saline perfusion, we transcardially perfused mice with 4% PFA (made in 0.1 M phosphate buffer, pH 7.4), removed their brains from the skull, and submerged them in 4% PFA overnight. 30 μm coronal slices were obtained from cryostat (Leica CM3050 S), serially following dehydration with a gradient sucrose solution (10-30% w/v). The mPFC tissue was collected using stereotaxic coordinates based on the Paxinos and Franklin mouse brain atlas, with anteroposterior (AP) coordinates ranging from +1.94 mm to +1.70 mm anterior to Bregma. These sections were used to assess the expression of synthetic enzymes: DAGL-α, enzymes related to glutamine– glutamate/GABA cycling: glutamine synthetase (GS) and GAD67, cannabinoid receptors: CB1 and CB2, inflammatory signalling: HMGB1 (High Mobility Group Box 1), NF-κB (Nuclear Factor kappa B), and TLR4 (Toll-Like Receptor 4), in the mPFC region of aged brain. Using a Nikon AX/AX R Confocal microscope, immunostained tissues (3 tissues per animal, total 12 tissues /group) were scanned to produce Z-stacks (1024 × 1024 pixels) were captured (60×) over the complete tissues at 1-μm intervals. To ensure uniformity and consistency, the data were exclusively collected from 20-μm-thick Z-stacks to minimize variability in data due to differences in mounted tissue thickness. Overall, the presented expression levels of DAGL-α, GS, GAD67, CB1, CB2, HMGB1, NF-κB and TLR4 were quantified based on channel intensity in Z-stacks (3D volume: 210 × 210 × 20 µm^3^), and subsequently converted to digital values representing the area of protein expression. A Nikon AX/AX R Confocal microscope at NIPER Hajipur produced the Z-stacks of three immunostained coronal slices per animal(Kesharwani et al., 2025a; Kesharwani et al., 2025b; Parihar and Limoli, 2013; Singh et al., 2025). ***Supplementary File 1*** contains detailed techniques and processes for immunofluorescence and confocal microscopy.

### Neurotransmitter estimations

mPFC tissues were homogenized in 400 µL of methanol and spiked with the internal standard, isoprenaline (100 ng/mL), followed by thorough vortexing. The homogenates were centrifuged at 12,000 rpm for 10 minutes, and the resulting supernatants were vacuum-dried for 2 hours. The dried residues were reconstituted in 200 µL of methanol, vigorously mixed, and centrifuged again at 12,000 rpm for 5 minutes. A 100 µL aliquot of the final supernatant was transferred into autosampler vials for quantification of glutamate, glutamine, and GABA using a Liquid Chromatography–High Resolution Mass Spectrometry (LC-HRMS) system at NIPER Hajipur(Kesharwani et al., 2025a; Kesharwani et al., 2025b). Detailed methodologies for neurotransmitter analysis are provided in ***Supplementary File 1***.

### Statistical Analyses

All statistical analyses were performed using GraphPad Prism 8.0.1. Data from the first group were analysed by one-way ANOVA followed by Tukey’s multiple comparisons test. Tukey’s multiple comparison was performed to determine significance differences among the groups. While for second group assigned for mitigating strategy are analysed by using unpaired t-test. Correlation analyses between DAGL-α activity and behavioral performance were performed using Spearman’s rank correlation coefficient (r). Correlations were interpreted as r values ranging from −1 to +1, indicating negative or positive associations, respectively, with r = 0 indicating no correlation. Data are presented as mean ± standard error of the mean (SEM). *p < 0.05; **p < 0.01; ***p < 0.001; and ****p < 0.0001.

## Results

### Aging drives progressive dysfunction in learning, mood and memory behaviour

We investigated the impact of aging on learning, memory, and behavioral functions in mice. Animals were subjected to the TO task ***(Fig. 2A)***. One-way ANOVA on time spent exploring object at phase 1 and phase 2 in TO task, revealed a significant effect of age on novelty preference (F_5, 54_ = 15.97, p < 0.0001). Further, Tukey’s multiple comparison test revealed that 20-Month aged mice had no preference for phase 1 object (p = 0.8564), while mice at age 4- and 12-Month showed significant (4-Month: p < 0.001, 12-Month: p = 0.0105) preference for the phase 1 object ***(Fig. 2B)***. Analysis of the DI by one-way ANOVA revealed a significant effect of age on preference of phase 1 (F_2, 27_ = 11.58, p = 0.0002). Further Tukey’s multiple comparison revealed that aging significantly impaired memory in aged mice, as evidenced by the lower DI in 12- and 20-Month mice as compare to 4-Month mice (4-Month vs 12-Month: p = 0.0476, 4-Months vs 20-Month: p = 0.0001; ***Fig. 2C***). These data demonstrate that aging potentially impairs memory performance in 12-Month and 20-Month aged mice. In addition, one way ANOVA analysis of total exploration time suggested no significant alteration between each group.

**Fig. 2:**
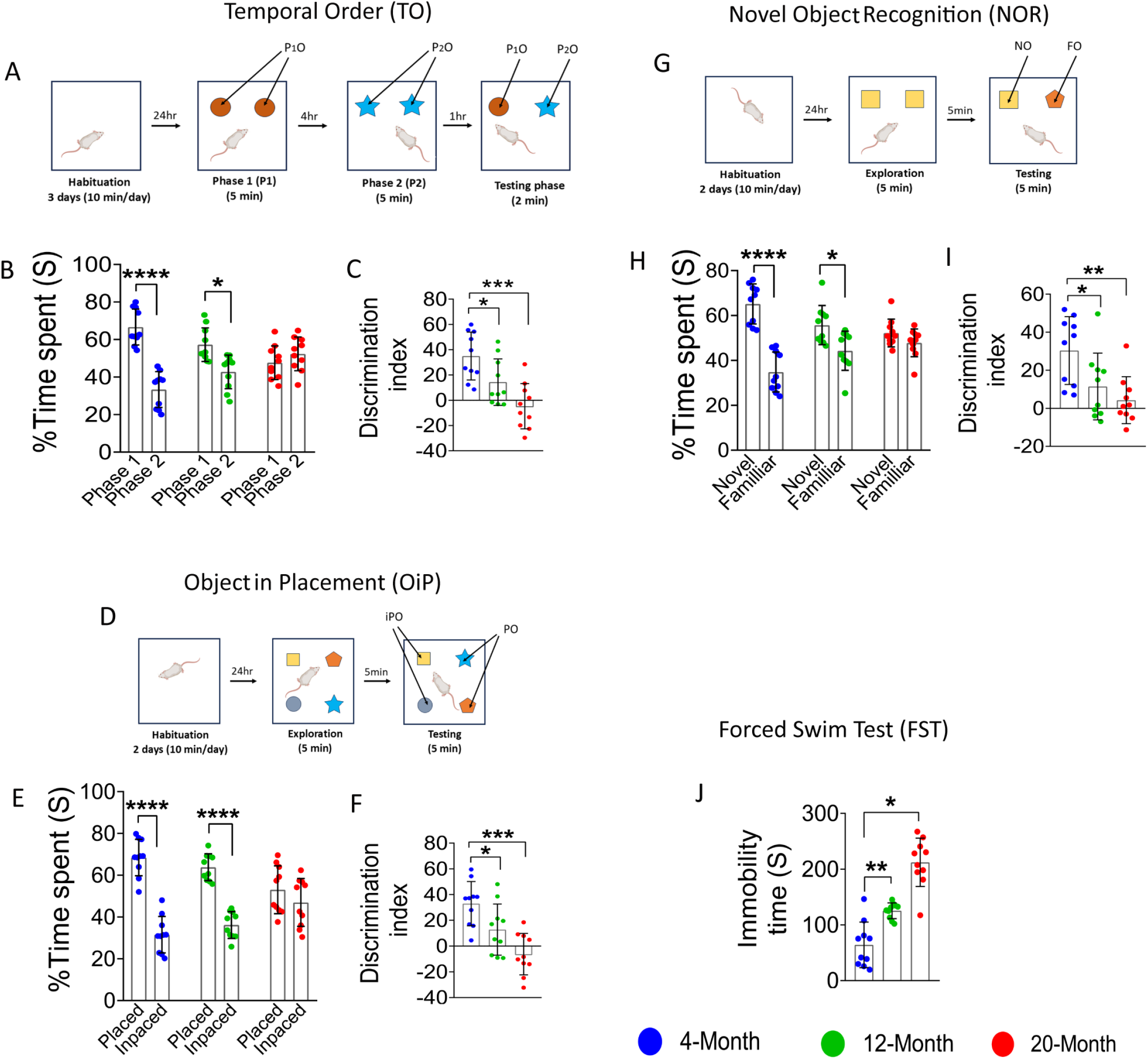
Aging induces cognitive impairment. TO: Schematic representation of the TO task (A). Aging significantly reduced the preference of Phase 1 object (B), and reduced DI (C). OiP: Schematic representation of the OiP task (D).Aged mice significantly reduced preference for placed object (E), approved by reduced DI (F). NOR: Schematic representation of the NOR task (G), Similarly, Preference of novelty was significantly reduced in aged mice (H), as indicated by DI (I). FST: aged mice displayed increased depressive-like behavior, as evidenced by, enhanced immobility time(J). Data are represented as mean ± SEM, one-way ANOVA followed by Tukey’s multiple comparison test. *p < 0.05, **p<0.01, ***p < 0.001, ****p < 0.0001, N = 10 per group. TO: Temporal Order, OiP: Object in Placement, NOR: Novel Object Recognition, DI: Discrimination Index, P1O: Phase 1 object, P2O: Phase 2 object, iPO: Inplaced object, PO: Placed object, NO: Novel object, FO: Familiar object.

Following TO task, we performed OiP test ***(Fig. 2D)***, one-way ANOVA analysis on time spent exploring at placed and inplaced object revealed a significant effect of age on novelty preference (F_5, 54_ = 26.12, p < 0.0001). Further, Tukey’s multiple comparison revealed that 20-Month aged mice had no preference for location (p = 0.6698), while mice at aged 4 and 12-Month showed significant (4-Month: p < 0.001, 12-Month: p < 0.0001) preference for the novel location ***(Fig. 2E)***. Analysis of the DI by one-way ANOVA revealed a significant effect of age on preference of novel location (F_2, 27_ = 12.20, p = 0.0002). Further Tukey’s multiple comparison revealed lower DI in 12- and 20-Month mice as compare to 4-Month mice (4-Month vs 12-Month: p = 0.0438, 4-Month vs 20-Month: p = 0.0001 ***(Fig. 2F)***. In addition, one way ANOVA analysis of total exploration time suggested no significant alteration between each group.

Following OiP task, we performed NOR test ***(Fig. 2G)***, one-way ANOVA analysis on time spent exploring at novel and familiar object revealed a significant effect of age on novelty preference (F_5, 54_ = 16.59, p < 0.0001). Further, Tukey multiple comparison revealed that 20-Month aged mice had no preference for novel object (p = 0.8266), while mice at aged 4- and 12-Month showed significant (4-Month: p < 0.001, 12-Month: p = 0.0278) preference for the novel object ***(Fig. 2H)***. Analysis of the DI by one-way ANOVA revealed a significant effect of age on preference of novel object (F_2, 27_ = 7.021, p = 0.0035). Further Tukey’s multiple comparison revealed lower DI in 12 and 20-Month mice as compare to 4-Month mice (4-Month vs 12-Month: p = 0.0361, 4-Month vs 20-Month: p = 0.0033 ***(Fig. 2I)***. One-way ANOVA analysis of total exploration time suggested no significant alteration between each group.

### Aging intensifies depressive like behavior in mice

The assessment of immobility duration in the FST indicated depressive-like behavior among the different mouse groups. The results of a one-way ANOVA demonstrated a significant increase in floating duration with advancing age (F_2, 27_ = 44.31, p < 0.0001; ***Fig. 2J)***. Further, Tukey’s multiple comparison indicated that mice at 12 and 20-Month exhibited a significant increase in floating time (4-Month vs 12-Month: 0.0017, p < 0.0001, 4-Month vs 20-Month: p < 0.0001) relative to those at 4-month mice.

### Aging alters eCB tone: Evidence from 2-AG deficiency in mice

ELISA kits measured the concentration of 2-AG in the mPFC. One-way ANOVA revealed a substantial reduction in 2-AG levels in mice (F_2, 27_ = 51.84, p < 0.0001). Further, Tukey’s multiple comparison demonstrated a lower quantity of 2-AG in mice at 12- and 20-Month compared to 4-Month. (4-Month vs 12-Month: p < 0.0001; 4-Month vs 20-Month: p < 0.0001; ***Fig. 3A***). The observed reduction at 12- and 20-Month reveals that age advancement is associated with a reduced amount of 2-AG in the mPFC.

**Fig. 3:**
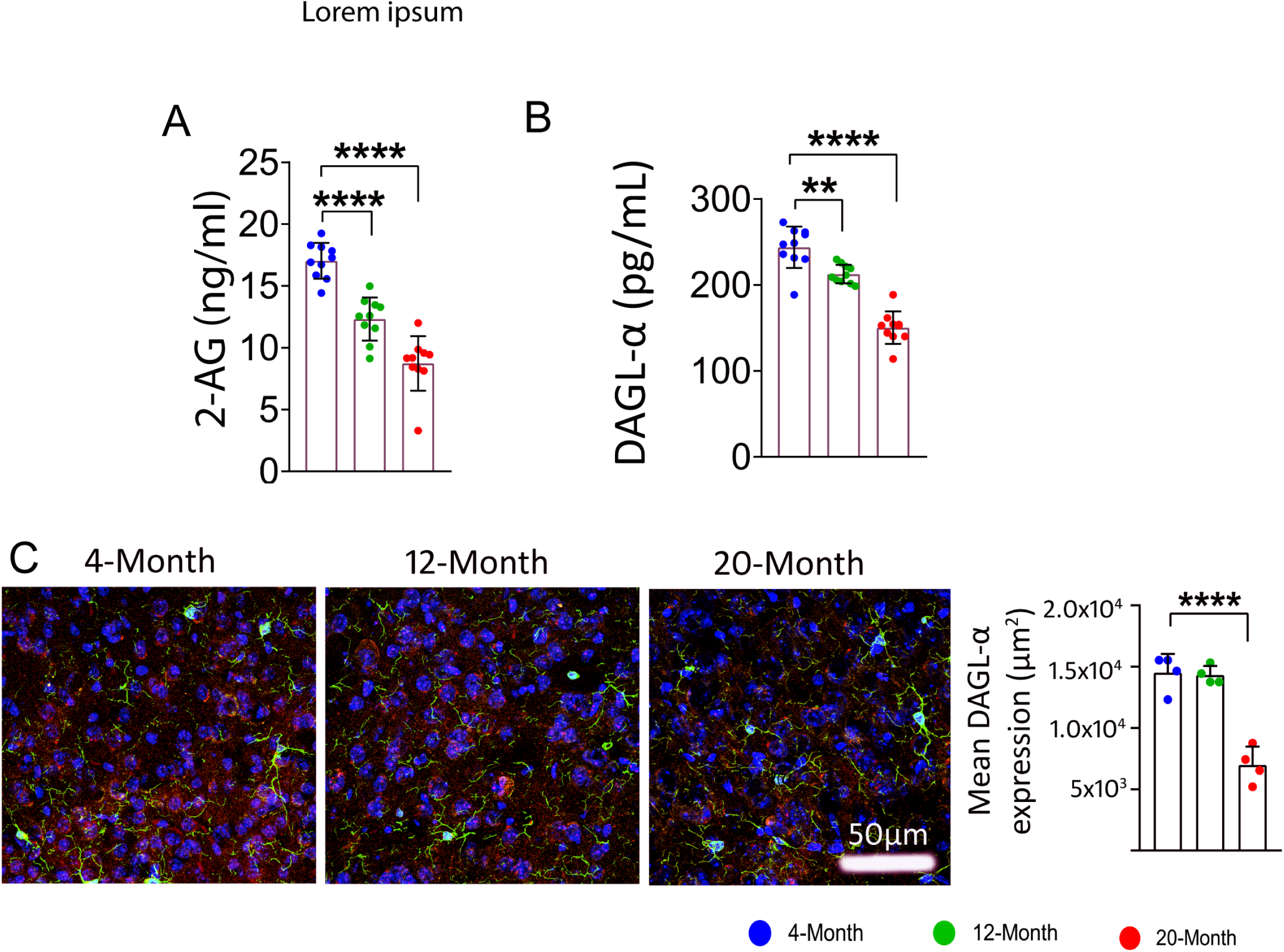
Temporary decrease in 2-AG levels in the mPFC of aged mice. Older mice exhibited lower levels of 2-AG **(A)** and diminished activity of DAGL-a **(B)**. Confocal images illustrate a decrease in DAGL-a expression associated with aging in the mPFC **(C)**. Green: microglia (IBA1 +ve), Red: DAGL-a, Blue: nuclear stain (DAPI +ve). Data are represented as mean ± SEM, one-way ANOVA followed by Tukey’s multiple comparison test. ***p < 0.001, ****p < 0.0001. N = 10 per group. 2-Ag: 2-Arachidonoyl Glycerol, DAGL-α: Diacylglycerol lipase-α.

### Impaired DAGL-α activity in the aged brain: A mechanistic link to eCB deficits

To investigate the specific mechanisms underlying the decreased levels of 2-AG in the mouse brain, we examined the expression and activity of diacylglycerol lipase-α (DAGL-α). Our results demonstrate that aging significantly alters DAGL-α activity (DAGL-α: F_2, 27_ = 64.52, p < 0.0001). Tukey’s multiple comparisons test further revealed a significant decrease in DAGL-α activity in mice aged 12-Months and 20-months compared to 4-month-old mice (4-Month vs. 12-month: p = 0.0026; 4-Month vs. 20-Month: p < 0.0001; ***Fig. 3B***). In addition, IF analysis confirmed these findings, showing altered expression of DAGL-α expression in aged mice (F_2, 9_ = 43.39, p < 0.0001). Further, Tukey’s multiple comparison test revealed lower expression of DAGL-α in 20-Month aged mice (p < 0.0001). Collectively, these results suggest that aging impairs DAGL-α activity and expression, leading to reduced 2-AG synthesis in the mouse brain. ***(Fig. 3C)***.

### 2-AG levels correlate with cognitive performance in mice

The process of aging leads to a reduction in 2-AG production and the synthesis of the DAGL-α enzyme. Consequently, we investigated the correlation between cognitive test performance and the alterations in 2-AG level and its synthesis enzymes as in each mouse. In our analysis, we did not observe a significant correlation in the groups of 4-Month and 12-Month mice (TO: 4-Month: r = 0.1472, p = 0.6849; 12-Month: r = 0.1400, p = 0.6996, NOR: 4-Month: r = −0.0945, p = 0.795; 12-Month: r = 0.4721, p = 0.1683). However, in the 20-Month group, we identified a significant correlation (TO: r = 0.7026, p = 0.008, NOR: r = 0.7026, p = 0.0235) between the concentration of 2-AG and performance on the DI-dependent task ***(Fig. 4A & C)***. Additionally, we examined the similar relationship between DAGL-α activity in each mouse and their performance on the DI dependent task. Our analysis revealed no significant correlation in the 4- and 12-Month aged mice (TO: 4-Month: r = 0.4541, p = 0.1874, 12-Month: r = 0.6312, p = 0.0503, NOR: r =0.3455, p = 0.3304).

**Fig. 4:**
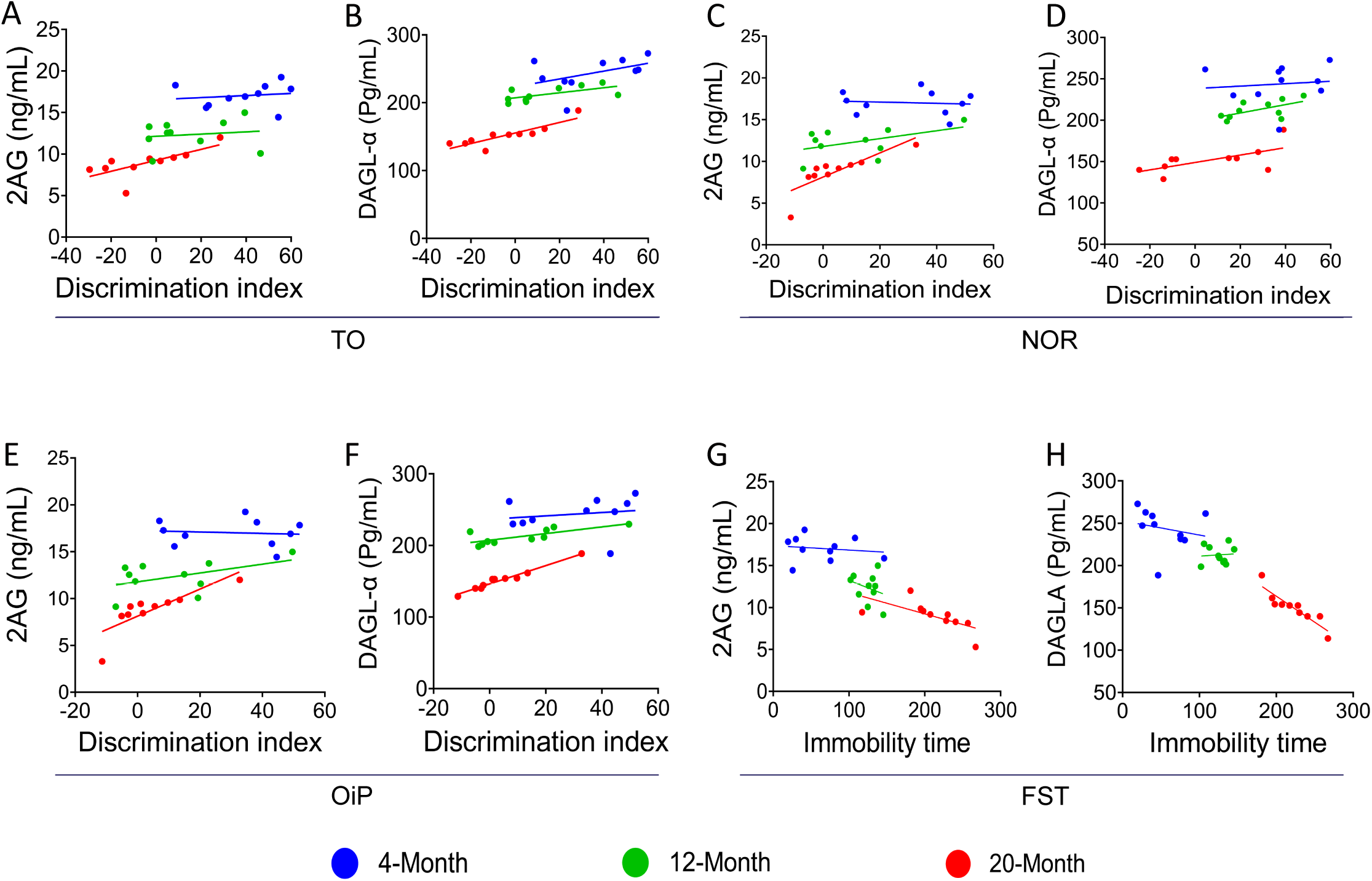
Correlation analysis between cognitive performance and eCB markers in aged mice. An analysis of Spearman’s correlation was conducted to evaluate the association between distinct cognitive performance metrics and the concentrations of 2-AG along with its synthesizing enzyme, DAGL-α. TO task: aging showed a potential correlation with impaired temporal memory and reduced level of 2-AG **(A)** and DAGL-α **(B)**. NOR task: aged mice exhibited a significant correlation between impaired recognition memory and decreased 2-AG **(C)** and DAGL-α **(D)** level. OiP task: no significant correlation was observed between object in place memory and 2-AG **(E)** or DAGL-α **(F)** levels. FST: levels of 2-AG **(G)** and DAGL-α **(H)** are significantly correlated with enhanced depressive-like behavior in aged mice. Data are presented as linear regression. **N** = 10 mice per group. 2-AG: 2-Arachidonoylglycerol; DAGL-α: Diacylglycerol lipase-alpha; TO: Temporal Order; NOR: Novel Object Recognition; OiP; Object in Placemnt, FST: Forced Swim Test

In contrast, the and 20-Month aged mice exhibited significant correlations between DAGL-α activity and DI dependent task performance (TO: 20-Month: r = 0.8690, p = 0.0011, NOR: 20-Month r = 0.9939, p < 0.0001; ***Fig. 4B & D)***. Furthermore, in OiP, there was no significant association between 2-AG or DAGL-α levels and cognitive performance. In our analysis, we did not observe a significant correlation in the groups of 4-Month, 12-Month and 20-Month mice (4-Month: r = −0.2731, p = 0.4452; 12-Month: r = 0.3186, p = 0.3697; 20-Month: r = 0.5739, p = 0.0828; ***Fig. 4E)***. As a result, memory for objects in the place is not reliant on mPFC 2-AG level at any age, and a reduction in it does not affect cognitive function. Additionally, we examined the similar relationship between DAGL-α activity in each mouse and their performance on the DI dependent task. Our analysis revealed no significant correlation in the 4-, 12- and 20 Month-aged mice (4-Month: r = 0.1059, p = 0.7709; 12-Month: r = 0.5971, p = 0.0684; 20-Month: r = 0.6374, p = 0.0474; ***Fig. 4F)***.

### 2-AG and the depressive like behaviour in mice

To assess whether the altered concentrations of 2-AG and DAGL-α were associated with depressive-like behavior in mice, we examined the levels of 2-AG and its enzyme, DAGL-α, in each mouse and the duration of their immobility. No significant correlation was observed between reduced 2-AG levels and increased depressive-like behavior in mice at 4- and 12-Month of age (4-Month: r = −0.1475, p = 0.6844; 12-Month: r = −0.2888, p = 0.4184). At 20-Month, a notable correlation emerged between reduced 2-AG levels and an increase in depressive-like behavior, characterized by extended durations of immobility (20-Month: r = −0.6495, p = 0.0421; ***Fig. 4G)***. Identical patterns were observed in DAGL-α. In mice, the analysis revealed no correlation between DAGL-α activity and immobility time at 4- or 12-month (4-Month: r = −0.2020, p = 0.5758; 12-Month: r = 0.07619, p = 0.8343). However, a significant correlation emerged at 20-month (20-Month: r = −0.9031, p = 0.0003; ***Fig. 4H)***.

### Age-related dysregulation of Glutamatergic and GABAergic balance in the mPFC

One-way ANOVA revealed a potential impact of aging on glutamine, glutamate, and GABA levels in the mPFC (glutamine: F_2, 12_ = 13.16, p = 0.0009; glutamate: F_2,12_ = 34.68, p < 0.0001; and GABA: F_2,12_ = 24.71, p < 0.0001). Further, Tukey’s multiple comparison revealed a decrease in glutamine and GABA levels in 20-Month aged mice (glutamine: p = 0.0007 and GABA: p < 0.0001) compare to 4-Month aged mice, similar analyses revealed that aging increased the levels of glutamate in 12- and 20-month aged mice (12-Month: p = 0.0145, 20-Month: p < 0.0001) ***(Fig. 5A, B & C)***. This data suggested that aging potentially altered glutamine-glutamate/GABA cycle in 20-month aged mice.

**Fig. 5:**
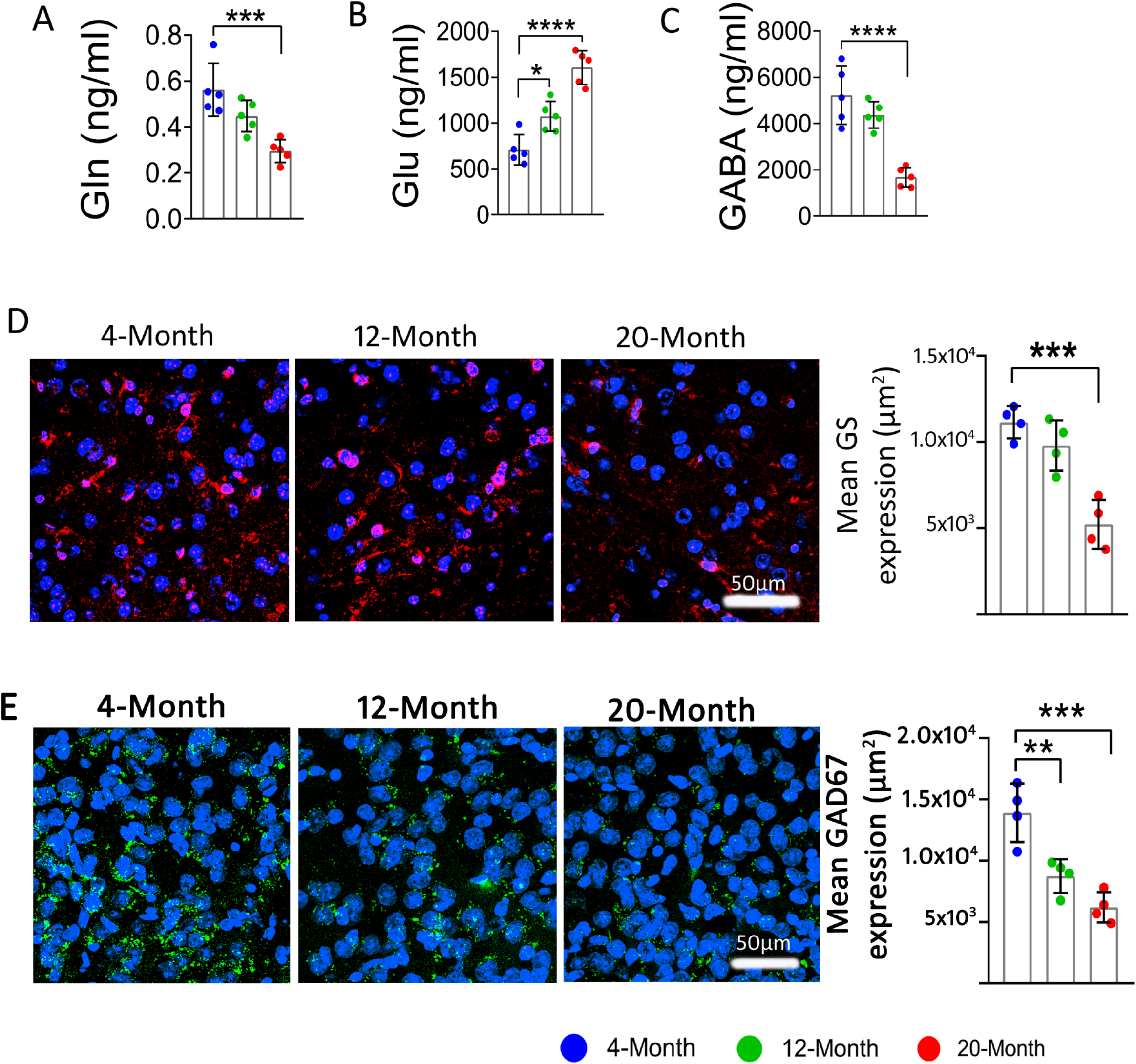
Aging altered glutamine-glutamate/GABA cycle in mPFC: Aged mice showed reduced level of glutamine **(A)**, glutamate **(B)**, and GABA **(C)**, Representative confocal images showing reduced expression of glutamine synthetase **(D)** and GAD67 **(E)** with advance aging in mPFC. Red: glutamine synthetase (GS +ve), Green: GAD67, Blue: nuclear stain (DAPI +ve). Data are represented as mean ± SEM, one-way ANOVA followed by Tukey’s multiple comparisons. ***p < 0.001, ****p < 0.0001. For IF: N = 4, For Neurotransmitter estimation N=5 per group. GS: glutamine synthetase, GAD67: Glutamate decarboxylase 67.

### Aging impairs glutamine–glutamate/GABA cycling by suppressing glutamine Synthetase and GAD67 expression in mPFC

To check the underlying mechanism of altered glutamine-glutamate/GABA cycle in aged mice brain, we have analysed the expression of the major enzymes, glutamine synthetase and GAD 67, which involved in the synthesis of glutamine and GABA. One-way ANOVA of Glutamine Synthetase ***(Fig. 5D)*** and GAD67 ***(Fig. 5E)*** suggested altered expression of GS and GAD67 in aged mice (GS: F_2, 9_ = 22.96, p = 0.0003; GAD67: F _2, 9_ = 20.34, p = 0.0005). Further Tukey’s multiple comparison test revealed that aging potentially reduce the expression of GS and GAD 67 in 20-month aged mice (GS: p = 0.0002 and GAD 67: p = 0.0003). This altered expression of GS and GAD67 in aged brain could be the possible mechanism behind the altered glutamine–glutamate/GABA.

### Age-Dependent modulation of CB1 and CB2 receptors in the mPFC

To determine the effect of aging on CB1 and CB2 receptors modulation in mPFC, the receptor density was examined by IF. One-way ANOVA of CB1 and CB2 receptors expression revealed a significant effect of aging (CB1: F_2, 9_ = 16.67, p = 0.0009, CB2: F_2, 9_ = 19.27, p = 0.0006) in aged mice brain (***Fig. 6A & B***).

**Fig. 6:**
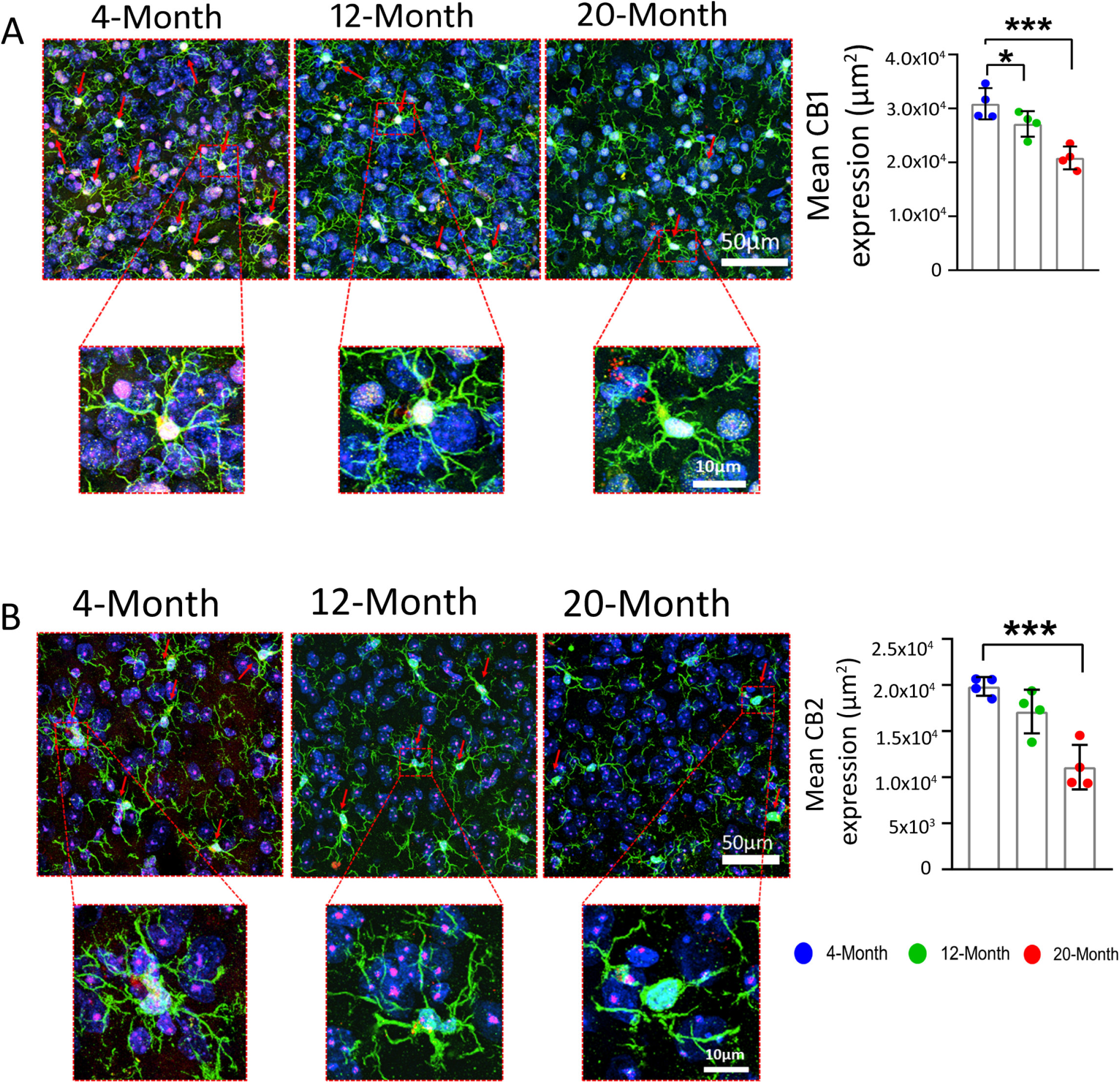
Age-related remodelling of microglial CB1/CB2 receptors in the mPFC. CB1 **(A)** and CB2 **(B)** receptor protein was significantly reduced in the mPFC of aged mice, while treatment with CBD effectively reversed these deficits. Representative confocal images show Green; microglia (IBA1+), Red: cannabinoid receptors (CB1 or CB2+ve), and Blue: nuclear staining (DAPI +ve). Data are represented as mean ± SEM, one-way ANOVA followed by Tukey’s multiple comparisons. *p < 0.05, ***p < 0.001. **N** = 4 mice per group.

Further, Tukey’s multiple comparison test revealed reduced expression of CB1 and CB2 receptor in the mPFC of 20-Month aged mice (CB1: p = 0.0006, CB2: p = 0.0004; ***Fig. 6A & B)***. These data suggest that CB1 and CB2 receptor-mediated endocannabinoid signalling in microglial cells contributes to the underlying mechanism.

### HMGB1-mediated TLR4/NF-κB pathway drives age-related microglial activation in the mPFC

To assess the effect of aging on HMGB1-mediated TLR4/NF-κB pathway and its link with microglial activation in aged mPFC. One-way Anova suggested that aging significantly promotes the HMGB1-mediated TLR4/NF-κB pathway in aged mice (HMGB1: F_2, 9_ = 24.24, p = 0.0002; TLR4: F_2, 9_ = 6.279, p =0.0196; NF-κB: F_2, 9_ = 10.95, p = 0.0039). Further, Tukey’s multiple comparison suggested a significant increase in neuroinflammation in the aged mouse (HMGB1: 4-Month vs 12-Month, p = 0.00360 and 4-Month vs 20-Month, p = 0.0001; TLR4: 4-Month vs 12-Month, p = 0.2100 and 4-Month vs 20-Month, p = 0.0001; NF-κB: 4-Month vs 12-Month, p = 0.0102, and 4-Month vs 20-Month, p = 0.0033). These findings suggest that aging may promotes key mediators of neuroinflammation in the aged brain. ***Fig. 7A, B, C***).

**Fig. 7:**
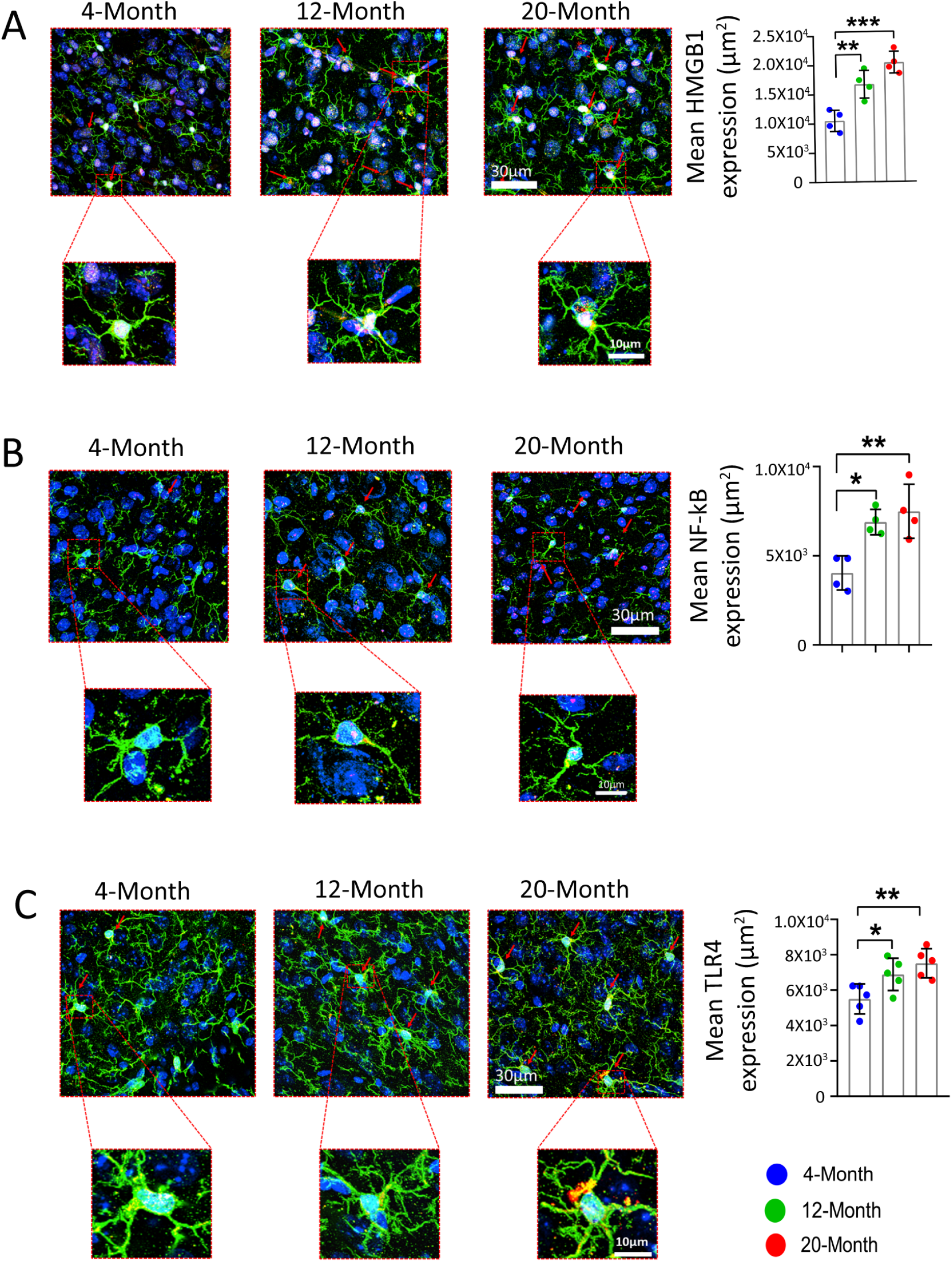
Aging promotes neuroinflammation in the mPFC. Representative confocal images showing elevated expression of inflammatory mediators HMGB1 (A), NF-KB (B), and TLR4 (C) in the aged mPFC, Green; microglia (IBA1+), Red: inflammatory mediators (HMGB1+ve, NF-KB +ve or TLR4 +ve), and Blue: nuclear staining (DAPI +ve). Quantification was performed using two-way ANOVA followed by Tukey’s multiple comparison test. *p < 0.05, **p < 0.01, ***p < 0.001. **N** = 4 mice per group.

### CBD a promising strategy to mitigate cognitive dysfunction by regulating altered eCB signalling

#### CBD improves mood and memory in aged mice

To check the effect of CBD on mood and memory, TO and NOR test have performed. One-way ANOVA analysis on time spent exploring objects at phase 1 and phase 2 revealed a significant effect of CBD on phase 1 object (F_3, 36_ = 10.35, p < 0.0001). Further, Tukey’s multiple comparison revealed that 20-Month aged mice had no preference for phase 1 object (p = 0.5552), while 20-Month mice treated with CBD showed significant preference for the phase 1 object (p < 0.0001; ***Fig. 8A)***. Analysis of the DI by unpaired t-test revealed a significant effect of CBD on DI (p = 0.0028; ***Fig. 8B)***. Unpaired t test revealed no significant changes in total object exploration time between each group.

**Fig. 8:**
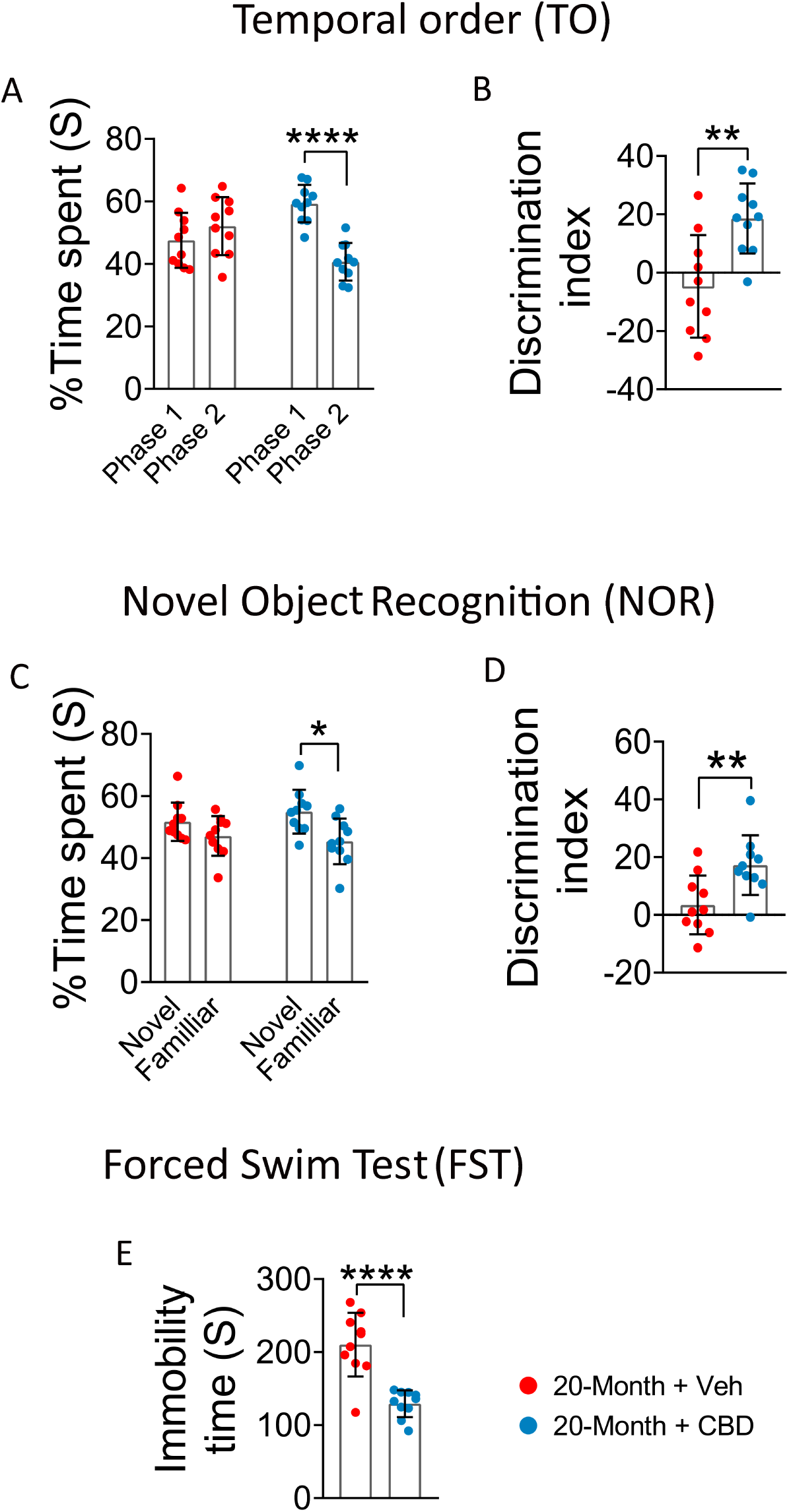
CBD improves age-related cognitive deficits across multiple behavioral paradigms. TO task: 20-Month + Veh group of mice exhibited a significant reduction in preference for the Phase 1 object, CBD treatment partially restored TO performance in aged mice **(A)**, which was further validated by a lower DI **(B)**. NOR task: Aged mice demonstrated a significant decrease in novelty preference, CBD treatment enhanced recognition memory in aged mice **(C)**, reflected in reduced DI **(D)**. FST: Aged mice showed increased depressive-like behavior, while CBD treatment significantly reversed these effects in aged mice **(E)**. Data are presented as mean ± SEM and analyzed using unpaired t-test. *p < 0.05, **p < 0.01, ****p < 0.0001. N = 10 mice per group. TO: Temporal Order; NOR: Novel Object Recognition; DI: Discrimination Index.

Following TO task, we performed NOR test, one-way ANOVA analysis on time spent exploring at novel and familiar object revealed a significant effect of CBD on novelty preference (F_3, 36_ = 4.131, p = 0.0129). Further, Tukey’s multiple comparison test revealed that 20-month aged mice had no preference for novel object (p = 0.442), while 20-Month mice treated with CBD showed significant preference for the novel object (p = 0.0161; ***Fig. 8C)***. Analysis of the DI by unpaired t-test revealed a significant effect of CBD on preference of novel object (p = 0.0076; ***Fig. 8D)***. These data demonstrate that CBD potentially regulates medial mPFC dependent memory in 20-Month aged mice. Unpaired t-test revealed no significant changes in total object exploration time between each group.

#### CBD potentially alleviates depressive-like behavior in aged mice

To check the impact of CBD on depressive like behavior of aged mice, our data were analysed by unpaired t-test, which suggested that CBD treatment significantly decreases (p < 0.0001) depressive like behavior in aged mice, evidenced by reduced immobility time ***(Fig. 8E)***.

#### CBD significantly regulates 2-AG and DAGL-α level in aged brain

To check the effect of CBD on eCB signalling, we have assessed the concentration of 2-AG in mPFC of aged brain. Our analysis through unpaired t-test revealed that CBD potentially enhanced the level of 2-AG (p < 0.0001) in 20-Month aged mice ***(Fig. 9A)***. Additionally, we have checked DAGL-α activity in mPFC, and unpaired t-test suggested CBD potentially promotes the activity of DAGL-α (p < 0.0001) enzyme in mPFC of 20-Month aged mice ***(Fig. 9B)***. Further, DAGL-α data was reconfirmed by the IF, unpaired t-test suggested that CBD potentially promotes the expression of synthesis enzyme in 20-Month aged mice (p = 0.0016; ***Fig. 9C***). These data revealed that CBD have potential activity to regulates cognitive dysfunctions by regulating the eCB signalling in aged mice.

**Fig. 9:**
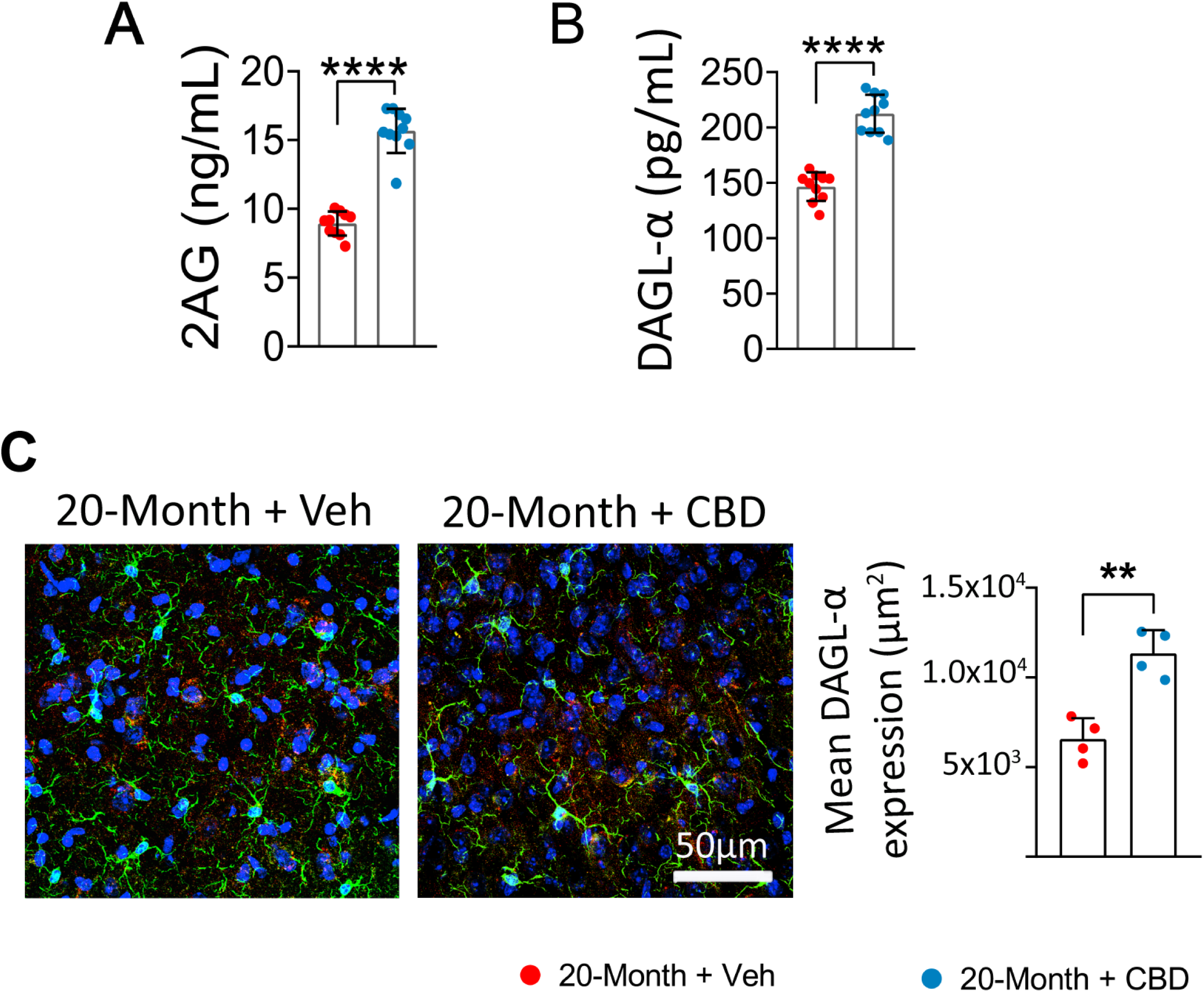
CBD restores 2-AG signalling in the aged mouse brain. Quantitative analysis showing that 20-Month + Veh group of mice exhibit significantly reduced levels of 2-AG, CBD treatment significantly restored it in mPFC (A), and decreased activity of its synthesizing enzyme DAGL-a mPFC, which was reversed by CBD (B). Representative confocal images illustrating reduced expression of DAGL-a in the mPFC of 20-Month + Veh group, reversed by CBD treatment (C), shows microglia Green: microglia (IBA1 +ve), Red: DAGL-a, Blue: nuclear stain (DAPI +ve). Data are presented as mean± SEM and analysed using unpaired t-test. **p < 0.01, ****p < 0.0001. N = 10 mice per group. 2-AG: 2-Arachidonoyl Glycerol; DAGL-a: Diacylglycerol lipase-alpha; CBD: Cannabidiol.

#### 2-AG and the cognitive performance of CBD treated aged mice

To check the effect of CBD on cognitive function, we have investigated the correlation between cognitive test performance and the alterations in 2-AG level and its enzymes in CBD treated mouse. In our analysis, we observe a significant correlation in the groups of 20-Month mice (TO: r = 0.7793, p = 0.0079, NOR: r = 0.8901, p = 0.0006). However, in the CBD treated 20-Month group, we did not identify a significant correlation (TO: r = −0.08547, p = 0.8144, NOR: r = 0.1303, p = 0.7197) between the concentration of 2-AG and performance on the DI-dependent task ***(Fig. 10A & C)***. Additionally, we examined the similar relationship between DAGL-α activity in each mouse and their performance on the DI dependent task. Our analysis revealed significant correlation in the 20-Month aged mice (TO: r = 0.6426, p = 0.0451, NOR: r = 0.7264, p = 0.0173). In contrast, the and CBD treated 20-month aged mice exhibited no significant correlations between DAGL-α activity and DI dependent task performance of CBD treated 20-month (TO: r = 0.5167, p = 0.1262, NOR: r = 0.4092, p = 0.2404; ***Fig. 10B & D)***.

**Fig. 10:**
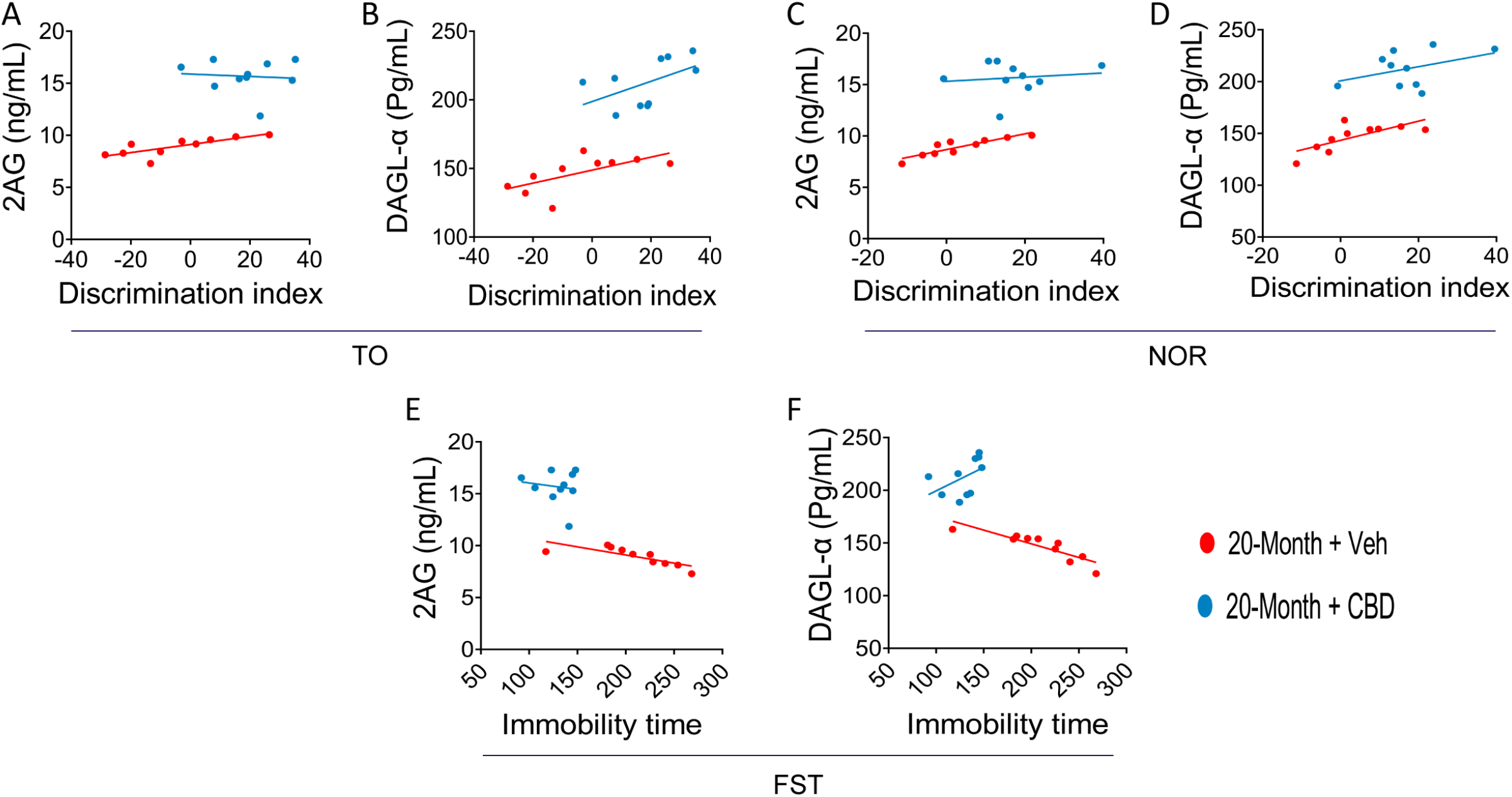
CBD modulates the correlation between cognitive behavior and eCB markers in aged mice. Spearman’s correlation analysis was performed to evaluate the relationship between individual behavioral performance and levels of 2-AG and its synthesizing enzyme DAGL-a in the mPFC of aged mice, with and without CBD treatment. TO task: Aging was associated with a negative correlation between cognitive performance and reduced levels of 2-AG **(A)** and DAGL-α **(B)**, which was attenuated in CBD-treated mice. NOR task: a significant correlation was observed between impaired recognition memory and decreased 2-AG **(C)** and DAGL-α **(D)** levels, while CBD treatment mitigating this relationship. FST: levels of 2-AG **(E)** and DAGL-α **(F)** showed a significant correlation with depressive-like behavior in aged mice, which was attenuated by CBD treatment. Data are presented as linear regression plots. N = 10 mice per group. 2-AG: 2-Arachidonoylglycerol; DAGL-α: Diacylglycerol lipase-alpha; TO: Temporal Order; NOR: Novel Object Recognition; FST: Forced Swim Test; CBD: Cannabidiol.

#### 2-AG and the depressive like behavior of CBD treated aged mice

To assess whether the regulated concentrations of 2-AG and DAGL-α by CBD treatment were manage depressive-like behavior in mice, we examined the levels of 2-AG and its enzyme, DAGL-α, in CBD treated aged mouse and the duration of their immobility. A significant correlation was observed between reduced 2-AG levels and increased depressive-like behavior in mice of 20-Months of age (r = −0.7809, p = 0.0077). while in 20-Month CBD treated mice, no significant correlation emerged between reduced 2-AG levels and an increase in depressive-like behavior, characterized by extended durations of immobility (r = −0.1444, p = 0.6907; ***Fig. 10E)***. Identical patterns were observed in DAGL-α. In mice, the analysis revealed significant correlation between DAGL-α activity and immobility time at 20-Month (r = −0.8729, p = 0.0010). However, no significant correlation emerged at 20-Month CBD treated mice (r = 0.4750, p = 0.1653; ***Fig. 10F)***.

#### CBD positively regulates the glutamine and glutamate level by modulating GS and GAD67 expression in mPFC of aged mice

To check the effect of CBD on glutamine-glutamate/GABA cycle, we have assessed their concentration in mPFC of aged mice. Our analysis through unpaired t-test revealed that CBD potentially increases level of glutamine in 20-Month aged mice (p = 0.0149; ***Fig. 11A)***, while CBD treatment significantly reduces glutamate level in 20-Month aged mice (p = 0.0011; ***Fig. 11B)***. However, we did not find any significant changes in GABA level of 20-Month aged mice (p = 0.0638; ***Fig. 11C)***. Additionally, to check the underlying mechanism of CBD on regulation of glutamine-glutamate/GABA cycle, we have examined the expression of GS and GAD67. Unpaired t-test revealed that CBD potentially promotes the expression of both of the enzymes (GS: p = 0.0001 and GAD67: p = 0.0449, ***Fig. 11D & E***). These finding indicated that CBD regulates glutamine-glutamate/GABA cycle by regulating the expression of two major enzymes involved in the synthesis of glutamine and GABA.

**Fig. 11:**
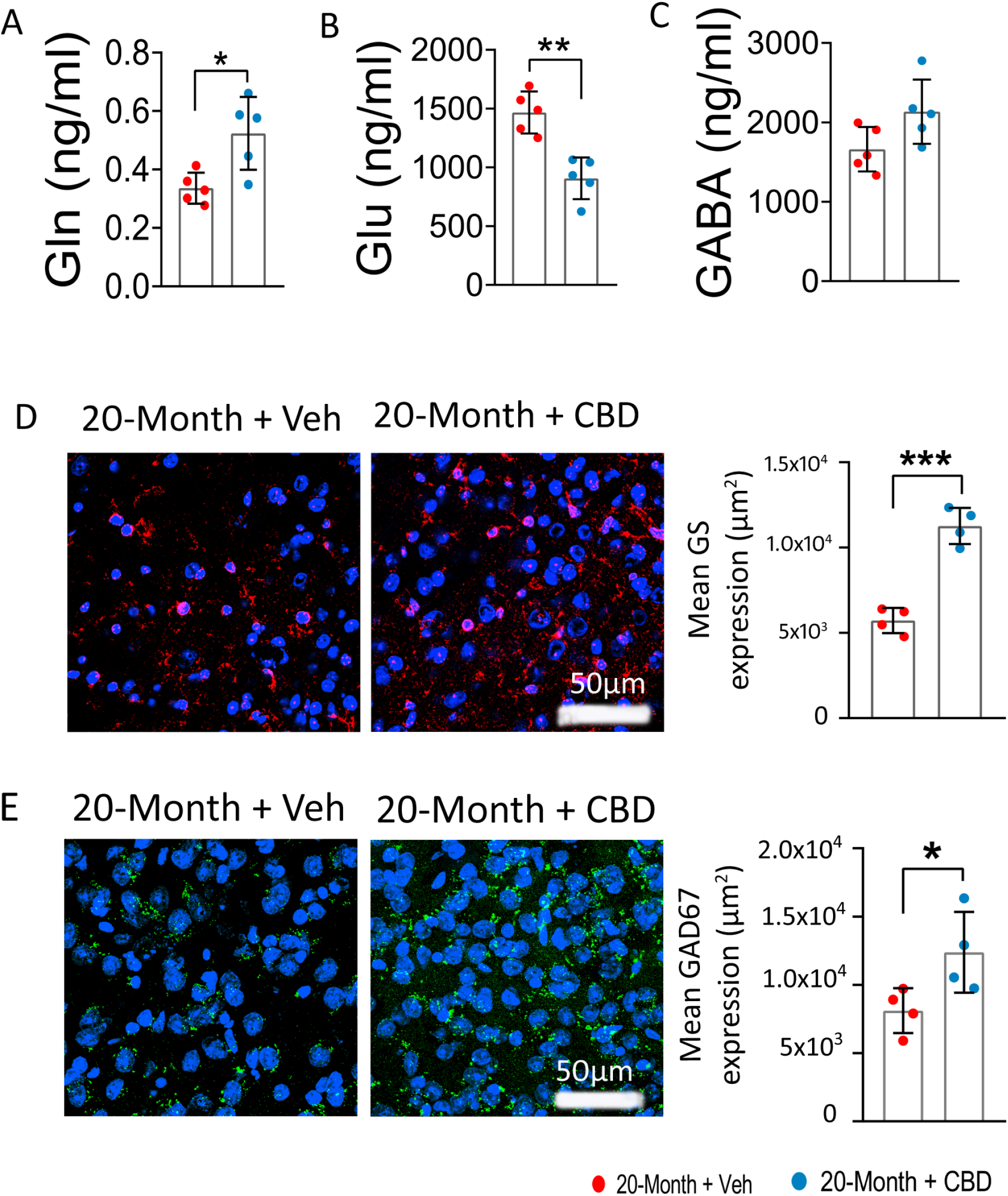
CBD reverses age-associated disruption in the glutamine-glutamate/GABA cycle in the mPFC. Quantitative analysis reveals that 20-Month + Veh group, exhibit significantly reduced levels of glutamine and glutamate, in the mPFC, which regulated by CBD treatment **(A-C)**, whereas no significant alteration in GABA level, N = 10 mice per group. Representative confocal images show reduced expression of glutamine synthetase **(D)** and glutamate decarboxylase 67 **(E)** in the mPFC of 20-Month + Veh group mice, both of which were markedly increased following CBD treatment. Red: glutamine synthetase (GS +ve), Green: GAD67, Blue: nuclear stain (DAPI +ve). Data are presented as mean ± SEM and analysed using unpaired t-test. *p < 0.05, **p < 0.01, ***p < 0.001. **N** = 4 mice per group. GS: Glutamine Synthetase; GAD67: Glutamate Decarboxylase 67; CBD: Cannabidiol.

#### CBD restores CB1 and CB2 receptor expression in the mPFC of aged mice

To assess the impact of CBD on CB1 and CB2 receptors in the mPFC of aged mice, receptor density was evaluated. Unpaired t-test revealed that CBD significantly enhanced the expression of CB1 and CB2 receptors in aged mice (CB1: p = 0.0121 and CB2: p = 0.0399 ***Fig. 12A & B***). These findings suggest that a potential mechanism underlying the benefits of CBD treatment involves modulation of eCB signalling in microglial cells.

**Fig. 12:**
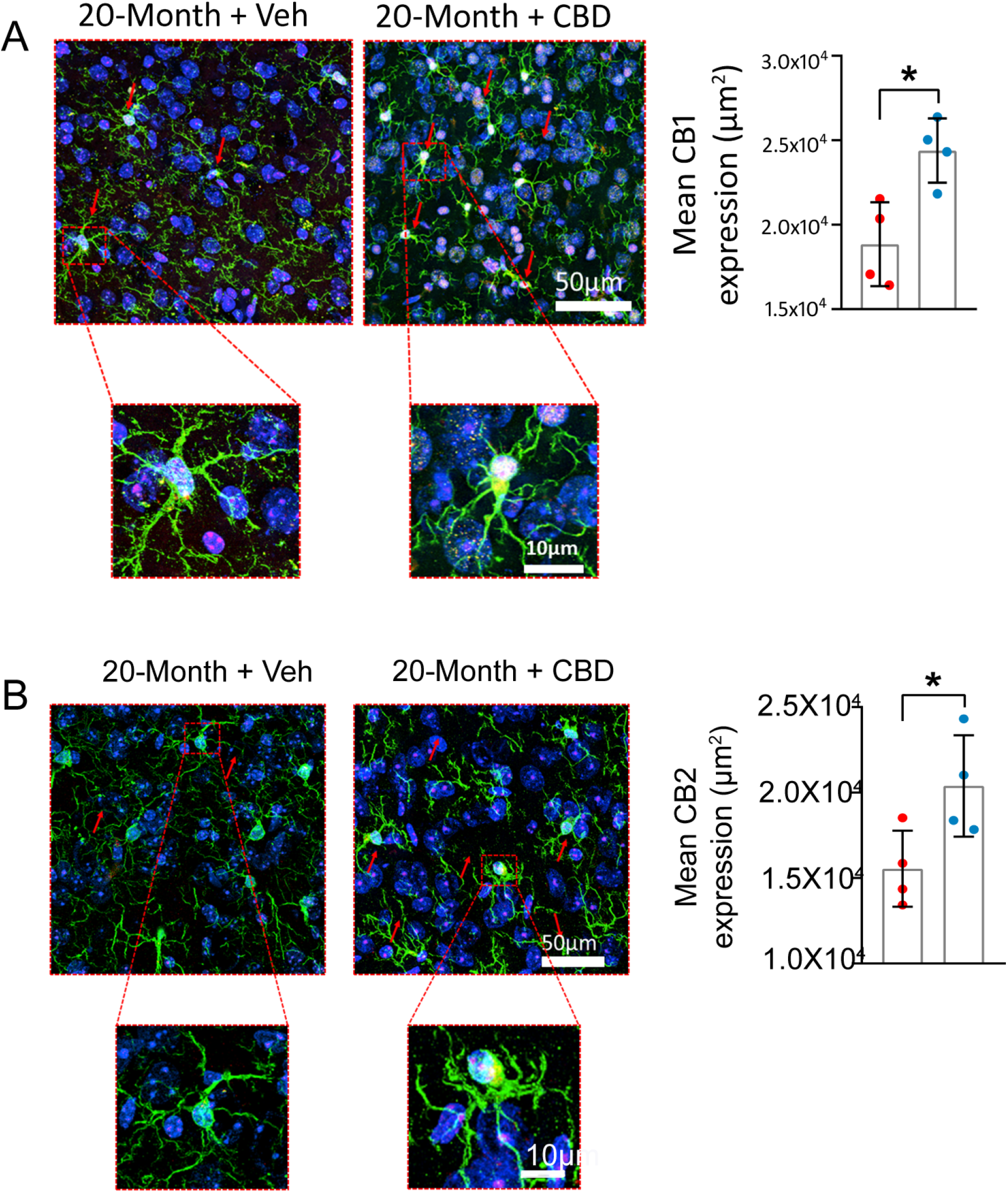
CBD restores CB1 and CB2 receptor expression on microglia in the mPFC of aged mice. Confocal images shows that 20-Month + Veh mice exhibit significantly reduced expression of cannabinoid receptors CB1 **(A)** and CB2 **(B)** in the mPFC. CBD treatment effectively reversed these age-related declines in receptor density. Representative confocal images illustrate Green; microglia (IBA1+), Red: cannabinoid receptors (CB1 or CB2 +ve), and Blue: nuclear staining (DAPI +ve). Data are presented as mean ± SEM and analysed using unpaired t-test. *p < 0.05. N = 4 mice per group. CB1: Cannabinoid Receptor 1; CB2: Cannabinoid Receptor 2; CBD: Cannabidiol.

#### CBD blocks HMGB1-TLR4/NF-κB driven microglial activation in the mPFC of aged mice

To assess the effect of CBD treatment on proinflammatory signalling, we quantified the number of HMGB1, TLR4 and NF-κB+ cells in the mPFC of aged mice. Unpaired t-test revealed that CBD significantly reduces the expression of proinflammatory mediators (HMGB1: p = 0.0015, TLR4: 0.0102 and NF-κB: p = 0.0415 ***Fig. 13A, B & C***). These findings indicate that CBD treatment may reduce critical drivers of neuroinflammation in the diabetic brain.

**Fig. 13:**
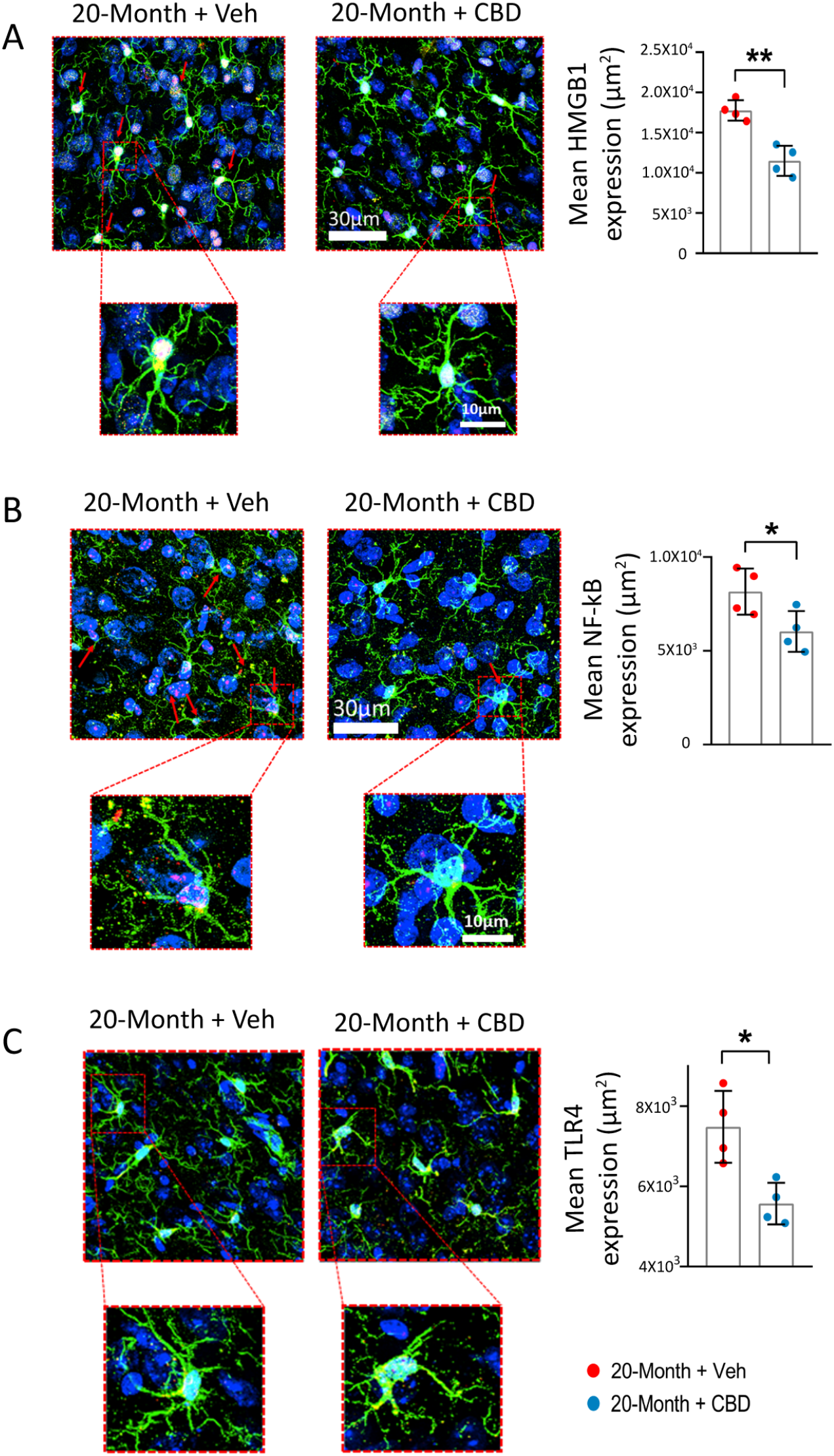
CBD attenuates age-induced neuroinflammation in the mPFC. Representative confocal images showing elevated expression of inflammatory mediators HMGB1 **(A)**, NF-KB **(B)**, and TLR4 (C) in the mPFC of 20-Month + Veh group. CBD treatment significantly reduced the expression of these inflammatory markers in aged mice. Green; microglia (IBA1+), Red: inflammatory mediators (HMGB1+ve, NF-KB +ve or TLR4 +ve), and Blue: nuclear staining (DAPI +ve). Data are presented as mean ± SEM and analysed using unpaired t-test. *p < 0.05, **p < 0.01. N = 4 mice per group. HMGB1: High Mobility Group Box 1; NF-KB: Nuclear Factor kappa B; TLR4: Toll-like Receptor 4; IBA1: Ionized calcium-binding adapter molecule 1; CBD: Cannabidiol.

## Discussion

Our study support that 2-AG plays a crucial role in cognitive aging, with its decline beginning in middle age and persisting with advancing age. Reduced 2-AG levels is associated with higher neuroinflammation, disrupted glutamine–glutamate/GABA cycle, cognitive deficits. Notably, we provide evidence that CBD, a non-psychoactive component of Cannabis sativa, improves mood and memory, reduces neuroinflammation and preserves cognitive function in aged mice. Furthermore, CBD enhances the expression of DAGL-α, a key enzyme responsible for 2-AG synthesis, suggesting a potential mechanism underlying its neuroprotective effects.

Cognitive aging was assessed using NOR, OiP, and TO tasks, revealing selective impairments in hippocampal- and prefrontal cortex-dependent spatial and temporal memory, while simple object recognition remained largely preserved. These findings are consistent with previous studies showing that aging specifically affects temporal memory, impairing the ability to distinguish past experiences based on temporal context (Kothari et al., 2017; Underwood and Thompson, 2016). To our knowledge, this is the first study to systematically link the temporal dynamics of cognitive decline with eCB. Interestingly, in a mouse model of diabetes, CBD treatment also significantly improved performance across various cognitive and depressive behavioral tasks including NOR and FST (Kesharwani et al., 2025a). Collectively, these findings support CBD as a promising intervention to mitigate age-associated cognitive decline in diverse models.

While our findings implicate eCB signalling in cognitive aging and neuroinflammation, the precise molecular mechanisms underlying the onset and progression of cognitive decline remain incompletely understood. Consistent with our observations (Fig. 5), previous studies have reported that cannabinoid receptors CB1 and CB2 are significantly reduced in aged mice compared to younger cohorts (Berrendero et al., 1998; Liu et al., 2003; Romero et al., 1998).

Genetic evidence further supports a critical role for CB1 receptors in maintaining cognitive function, with their loss accelerating cognitive decline and structural brain alterations (Bilkei-Gorzo et al., 2005; Saravia et al., 2017). Interestingly, a study in an Huntington disease (HD) model showed that the selective loss of CB1 receptors is mechanistically linked to GABAergic deficiency and the progression of cognitive decline (Saravia et al., 2017). Consistent with this, our study found CBD administration increases the expression of GAD67, an enzyme responsible for GABA synthesis (Fig 11). These findings suggest that CBD may improve cognitive deficits by regulating CB1 receptor signalling and GABA levels in aging, and may serve as a potential therapeutic strategy for HD. Importantly, decreased CB1 signalling has been shown to impair microglial phagocytosis, thereby limiting the clearance of cellular debris and exacerbating neuroinflammation (Stella, 2009). In line with this, our data demonstrate a strong association between altered eCB signalling and increased inflammatory responses (Fig. 6). Specifically, aging was associated with activation of the HMGB1–NF-κB–TLR4 signalling pathway, providing a mechanistic link between eCB dysregulation and neuroinflammation. Notably, CBD treatment restored the expression of eCB receptors and attenuated pro-inflammatory signalling in aged mice (Fig. 13), highlighting its therapeutic potential in mitigating cognitive aging and neuroinflammatory processes.

eCB signaling is closely linked to age-related disorders such as AD. Disruption of eCB signaling impairs the excitatory–inhibitory balance and synaptic homeostasis, leading to exacerbated behavioral deficits, particularly under conditions such as social isolation (Li, 2025; Melis et al., 2014; Zhang et al., 2009). These studies support a critical role for eCB signaling in regulating synaptic function in AD. Consistent with this, we found that CBD treatment is beneficial under conditions associated with cognitive dysfunction in normal mice, supporting its potential as a therapeutic agent for AD and associated dementias. Importantly, CBD increased the expression of DAGL-α and CB1/CB2 receptors in aged mice, suggesting that restoration of 2-AG biosynthesis may underlie its neuroprotective effects, including the prevention of memory deficits and neuroinflammation. Thus, our study indicates that targeting eCB signaling remains an attractive strategy for mitigating cognitive aging. In addition, our findings highlight another potential therapeutic avenue, selective enhancement of 2-AG levels. This could be achieved in two ways: by increasing the activity of DAGL-α, which promotes 2-AG synthesis, or by inhibiting monoacylglycerol lipase (MAGL), the enzyme that degrades 2-AG. While the former may require gene therapy approaches that are not readily accessible to all patients, MAGL inhibition represents a more feasible alternative that warrants further investigation.

In summary, our study suggests that targeting the eCB pathway represents a promising therapeutic strategy to improve cognitive deficits associated with normal aging and neurodegenerative disorders such as HD and AD.

## Abbreviations

eCB: endocannabinoid
CBD: Cannabidiol
CB1: type 1 cannabinoid receptor
CB2: type 1 cannabinoid receptor
CNS: central nervous system
DAGL-α: Diacylglycerol lipase-alpha
2-AG: 2-arachidonylglycerol
TO: Temporal Order
NOR: Novel Object Recognition
OiP: Object in Placement
FST: Forced Swim Test
GS: Glutamine synthetase
GAD67: Glutamate decarboxylase 67.

## Authors’ contributions

**AK**: Conceptualization, Data curation, Formal analysis, Methodology, Software, Validation, Visualization, Writing – original draft, Writing – review & editing. **DL:** Methadology, Formal analysis, Writing – review & editing. **AS:** Validation, Visualization, Writing – review & editing. **MK:** Validation, Visualization, Writing – review & editing. Validation, Visualization, Writing – review & editing. **VR**: Funding acquisition, Investigation, Project administration, Resources, Supervision. **VKP:** Conceptualization, Funding acquisition, Investigation, Project administration, Resources, Supervision, Validation, Writing – review & editing.

## Ethical approval

The research protocol was approved by the “Institutional Animal Ethics Committee (IAEC) at the National Institute of Pharmaceutical Education and Research in Hajipur, India under the approval number (Approval No: NIPER-H/IAEC/21/22), following the recommendations of Committee for the Control and Supervision of Experiments on Animals (CCSEA), Ministry of Environment and Forests, Government of India, New Delhi.

## Acknowledgement

We thank the Confocal-STORM Microscopy Facility at NIPER-Hajipur for microscopy support, and Mrs. Shreetoma Kundu, NIPER-Kolkata, for her assistance in capturing confocal images.

## Availability of data and material

Data generated in this study are included in the manuscript and supplementary information.

