## Supplementary file 1 for "Cannabidiol rescues age-associated cognitive decline in mouse model"

### ^d^Delhi Pharmaceutical Sciences and Research University, New Delhi - 110017, India

^e^Department of Regulatory Toxicology, National Institute of Pharmaceutical Education and Research, Hajipur - 844102, India

^#^Current affiliation: Florida Atlantic University, Stiles-Nicholson Brain Institute, Jupiter, Florida- 33458, USA

^@^Current affiliation: Central University of Haryana, Mahendergarh, Haryana-123031, India

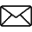
 **Corresponding Author**

Dr. Vipan K. Parihar,

Department of Pharmacology and Toxicology,

Department of Regulatory Toxicology

National Institute of Pharmaceutical Education and Research

Hajipur - 844102, Bihar, India

**Behavioral assessment:**

**Temporal Order (TO) test:** Following a three-day habituation period (10 minutes per day) in the open arena, a temporal order memory test was conducted, consisting of three distinct phases, each with slight modifications. In the first phase, mice were exposed to two identical objects for 5 minutes. They were then returned to their home cages for a 4-hour interval. During this break, both the arena and objects were sanitized using 70% alcohol. In the second phase, the mice were reintroduced to the arena containing two new objects of the same shape as those in phase one, again for 5 minutes. Subsequently, the animals were placed back in their cages for an additional hour in the same room, with the arena and objects once again cleaned using 70% ethanol. In the final phase, the mice were presented with one object from each of the previous two phases and allowed to explore for 5 minutes. An observer, blinded to the treatment groups, recorded the behavior, and results were analyzed using the discrimination index. Scoring was performed in accordance with established criteria for assessing temporal order memory (Acharya et al., 2016; Barker et al., 2007; Kesharwani et al., 2025b; Parihar et al., 2015).

**Novel Object Recognition (NOR) test:** After three days of habituation in an open field (10 minutes per day), animals underwent the Novel Object Recognition (NOR) test. Objects used across all behavioral tests were similar in material but varied in shape and color. To prevent displacement by the mice, objects were magnetized and positioned 7 cm from opposing corners and 16 cm apart. During the familiarization phase, mice were exposed to two identical objects for five minutes. Following this, they were returned to their home cages for a 5-minute interval, during which the arena and objects were sanitized with 70% ethanol and one object was replaced with a novel one of a different shape. The mice were then given another five-minute exploration period with both the familiar and novel objects. Object placement was counterbalanced across groups, and no object was reused in later tests. An observer blind to the treatment groups scored the trials. Exploratory behavior was recorded when a mouse approached within 1 cm of an object and directed its gaze toward it. A discrimination index was calculated to assess novelty preference or indifference (Acharya et al., 2016; Acharya et al., 2015; Kesharwani et al., 2025a; Kesharwani et al., 2025b; Parihar et al., 2015).

**Object in Place (OiP) task:** Following the NOR test, mice underwent a two-day habituation period in an open arena (10 minutes per day). On the third day, they were exposed to four distinct objects differing in shape, color, and size, for 5 minutes to allow familiarization. After this session, mice were returned to their home cages for 5 minutes in the same room. During this interval, the arena and objects were cleaned with 70% ethanol, and the positions of two of the four objects were swapped. All objects were presented equally to all groups to eliminate any bias toward a particular item. Mice were then reintroduced to the arena for another 5 minutes to explore the new object arrangement. Behavioral scoring was conducted by an observer blinded to group assignments, and performance was analyzed using the discrimination index (Acharya et al., 2016; Acharya et al., 2015; Barker et al., 2007; Kesharwani et al., 2025a; Kesharwani et al., 2025b).

**Forced Swim Test (FST):** The Forced Swim Test (FST) is a widely used behavioral assay to assess helplessness or depression-like behavior in rodents. In this test, rodents are placed in a water-filled cylinder, and their mobility is observed over a duration of 5 to 8 minutes. The period during which the animal remains mostly still, keeping its head above water without active escape attempts, is termed "floating behavior" and serves as an indicator of depression-like behavior. In this experiment, each mouse was placed in a glass beaker with an inner diameter of 10 cm and a height of 14.5 cm, filled with tap water at 18–20°C to a depth of 9.5 cm. Prior to testing, it was ensured that the water level prevented the animals from reaching the bottom with their hind limbs or tails. Each session lasted 5 minutes, during which immobility and struggling behaviors were recorded minute by minute. Struggling was defined as vigorous activity aimed at escaping the beaker, while immobility or floating was characterized by minimal movement sufficient to keep the head above water. After the session, each mouse was removed, gently dried with a towel or dryer, and returned to its home cage. The total immobility duration per mouse was calculated and used as a measure of depression-like behaviour (Acharya et al., 2015; Kesharwani et al., 2025a; Kesharwani et al., 2025b; Parihar and Limoli, 2013).

**Tissue processing for ELISA, immunofluorescence, and neurotransmitter estimation: Unlocking the molecular insights behind altered eCB signaling:** Following the completion of behavioral testing, comprehensive biological and immunohistochemical analyses were conducted to investigate the underlying molecular and cellular changes. The mice were allocated into two distinct cohorts. The first group underwent transcardial perfusion with 4% paraformaldehyde (PFA) followed by saline, and their brains were preserved for immunohistochemical evaluations. In contrast, the second group was perfused with saline alone, and their brain tissues were collected for ELISA-based assays and neurotransmitter quantification. The subsequent section details the methodologies employed, including tissue preparation, staining protocols, antibody specifications, and analytical procedures used to assess relevant biomarkers and cellular alterations (Kesharwani et al., 2025a; Kesharwani et al., 2025b; Singh et al., 2025).

**ELISA-based detection of proteins:** Quantitative analysis of 2-Arachidonoylglycerol (2-AG) and Diacylglycerol lipase-α (DAGL-α) levels in the mPFC tissue homogenates was performed using commercially available ELISA kits.

- **2-Arachidonoylglycerol (2-AG):** To ensure data accuracy and reproducibility, it is highly recommended that all samples and standards be analyzed in duplicate or triplicate. Each assay should also include a standard curve for precise quantification of target analytes. The procedure begins by adding 50 µL of Standard Diluent to the designated blank wells to establish a baseline. Subsequently, 50 µL of each prepared standard and sample is added to the appropriate wells. Following this, 50 µL of the Biotinylated Primary Antibody Working Solution is introduced into each well to facilitate antigen-antibody binding. The microplate is then sealed and incubated at 37°C for 60 minutes. After incubation, the wells are aspirated and washed four times with 1X diluted Wash Buffer to remove unbound substances. To ensure complete removal of residual buffer, the plate is inverted and tapped firmly on absorbent paper, and any remaining liquid on the bottom surface is wiped off to prevent interference with absorbance readings. Next, 100 µL of the Streptavidin-Horseradish Peroxidase (HRP) Conjugate Working Solution is added to each well. The plate is gently mixed to ensure even distribution, resealed, and incubated again at 37°C for 60 minutes. Following this second incubation, the washing step is repeated as previously described. Then, 100 µL of TMB substrate solution is added to all wells, and the plate is incubated at 37°C for 10 minutes without agitation to avoid elevated background and ensure optimal sensitivity. To terminate the enzymatic reaction, 100 µL of Stop Solution is added to each well, resulting in a color shift from blue to yellow. The absorbance is then measured at 450 nm using a microplate reader within 10–15 minutes of stopping the reaction to guarantee accurate quantification (Kesharwani et al., 2025a).
- **Diacylglycerol lipase (DAGL-α):** Designate seven wells for the standard curve and one well as the blank control. Add 100 µL of either the prepared standard working solutions or samples to the appropriate wells. Cover the plate and incubate at 37 °C for 80 minutes to allow antigen binding. Following incubation, discard the contents and wash each well three times with 200 µL of 1× Wash Buffer, allowing the buffer to sit in the wells for 1–2 minutes during each wash. After the final wash, blot the plate dry on absorbent paper. Next, add 100 µL of the Biotinylated Antibody Working Solution to each well. Reseal the plate and incubate at 37 °C for 50 minutes. Afterward, wash the wells three times using the same procedure. Then, dispense 100 µL of the Streptavidin-HRP Working Solution into each well, cover the plate, and incubate at 37 °C for another 50 minutes. This is followed by five wash steps to ensure the removal of unbound reagents. Subsequently, add 90 µL of TMB Substrate Solution to each well and incubate the plate in the dark at 37 °C for 20 minutes to allow color development. After this step, add 50 µL of Stop Reagent to each well. Gently tap the plate to mix the contents thoroughly. Finally, wipe the bottom of the plate to remove any condensation or residue and immediately read the absorbance at 450 nm using a microplate reader (Kesharwani et al., 2025a).

**PFA perfusion, Tissue Immunolabeling and Confocal Microscopy:** On the day following the final behavioral assessments, animals were euthanized, and their brains were processed for immunofluorescence evaluation to examine specific molecular markers in the mPFC. Mice were deeply anesthetized in a plexiglass induction chamber using isoflurane until complete cessation of respiration was observed, ensuring a surgical level of anesthesia. Once unresponsive, the thoracic cavity was carefully opened to expose the heart. An incision was made in the right atrium to facilitate drainage, and the left ventricle was cannulated to initiate transcardial perfusion. Perfusion was conducted in two phases: initially, 25 mL of ice-cold normal saline was perfused at a constant flow rate of 14–16 mL/min to flush out the blood, followed by 75 mL of 4% paraformaldehyde (PFA) in phosphate buffer (pH 7.4) to fix the tissues. This process ensured both vascular clearance and effective fixation. Following perfusion, brains were carefully dissected, post-fixed in 4% PFA overnight at 4 °C, then rinsed with phosphate buffer. To achieve cryoprotection, the tissues were successively immersed in ascending concentrations of sucrose solutions (10%, 20%, and 30% w/v), remaining in each until the brains sank, indicating full infiltration. Cryoprotected brains were embedded in cryostat embedding medium and sectioned coronally at a thickness of 30 µm using a cryostat (Leica CM3050 S), proceeding from posterior to anterior brain regions. Serial sections were collected and stored in 24-well plates containing PBS supplemented with 0.2% sodium azide to prevent microbial growth until further immunohistochemical analysis (Kesharwani et al., 2025a; Singh et al., 2025).

Three serial sections from each mPFC tissue sample were collected using stereotaxic coordinates based on the Paxinos and Franklin mouse brain atlas, with anteroposterior (AP) coordinates ranging from +1.94 mm to +1.70 mm anterior to Bregma ***(Supplementary fig. S1),*** to examine the expression of microglia, cannabinoid receptors (CB1 and CB2), enzymes involved in endocannabinoid (eCB) synthesis (DAGL-α), markers of glutamate/GABA cycling (glutamine synthetase and GAD67), and inflammatory cytokines (HMGB1, NF-κB, and TLR4) (Kesharwani et al., 2025a; Parihar et al., 2018, 2016). Tissue sections were first blocked with 10% normal goat serum (NGS) and 0.1% Triton X-100 (TTX-100) for 30 minutes at room temperature. Primary antibodies (IBA-1: rabbit polyclonal, G-Biosciences, 1:1000; glutamine synthetase: mouse monoclonal, CiteAb, MA5-27749; GAD67: rabbit polyclonal, Thermo Fisher, PA5-21397) were then applied in 2% NGS with 0.1% TTX-100 and incubated overnight at 4°C. After washing with PBS (three times, five minutes each), the sections were incubated for one hour at room temperature with appropriate secondary antibodies (Alexa Fluor 488 goat anti-rabbit for IBA-1 and GAD67; Alexa Fluor 594 goat anti-mouse for glutamine synthetase, Thermo Fisher, A32742) in 2% NGS. For double immunostaining, sections were also incubated overnight at 4°C with primary antibodies against CB1 (rabbit polyclonal, Thermo Fisher, 1:500, PA5-85080), CB2 (rabbit polyclonal, Thermo Fisher, 1:500, CB2-201AP), DAGL-α (rabbit polyclonal, Thermo Fisher, 1:1000, PA5- 23765), HMGB1 (rabbit polyclonal, Thermo Fisher, 1:1000, PA5-27378), NF-κB (rabbit polyclonal, Thermo Fisher, 1:500, PA5-27617), and TLR4 (mouse polyclonal, Abcam, 1:1000, ab22048), diluted in 2% NGS with 0.1% Triton X-100. After three PBS washes, CB1, CB2, DAGL-α, HMGB1, and NF-κB were visualized using Alexa Fluor 594 goat anti-rabbit secondary antibody (Thermo Fisher, 1:1000, A32740), while TLR4 was detected with Alexa Fluor 594 goat anti-mouse antibody (Thermo Fisher, 1:1000, A32742). Sections were incubated for one hour at room temperature with the secondary antibodies. Following antibody staining, tissues were counterstained with DAPI (Abcam, ab228549) for seven minutes, mounted in an antifade medium, and stored for imaging. Imaging was performed using a Nikon AX/AX R confocal microscope at NIPER-Hajipur. Z-stack images were collected from three sections per animal (12 sections per group) using 60× magnification at 1-μm intervals (1024 × 1024 pixels). Imaging parameters were kept consistent across all samples. Only staining patterns that met established criteria for specificity and intensity were included in quantitative analyses. Microglial expression of CB1, CB2, DAGL-α, HMGB1, NF-κB, TLR4, glutamine synthetase, and GAD67 was analyzed using Z-stack images. Prior to quantification, confocal Z-stacks were deconvoluted to correct for z-axis distortion. Immunoreactivity was quantified by measuring channel color intensities in the Z-stacks across the full tissue depth.

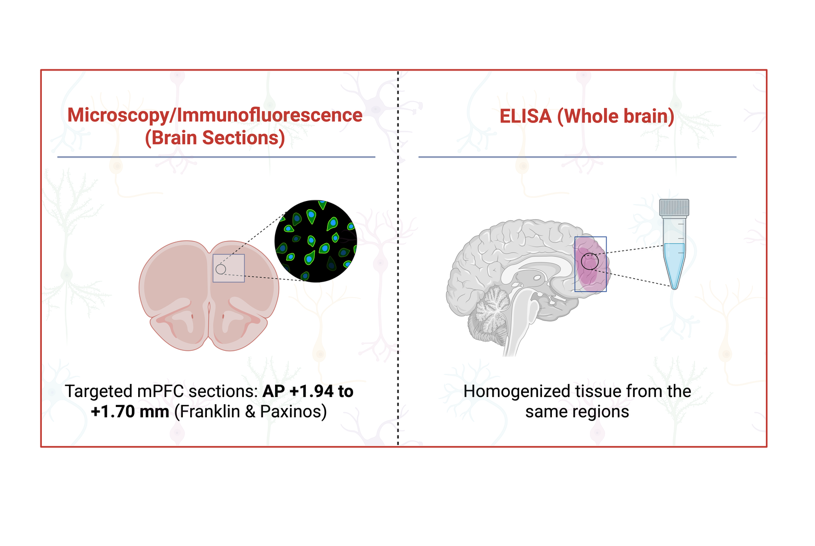

**Fig. S1:** Schematic showing mPFC analysis sites used for immunofluorescence and ELISA.

**Brain tissue processing for LC-HR/MS-based neurotransmitter profiling:** The mPFC, weighing approximately 50 mg, was carefully dissected on ice to preserve tissue integrity and prevent enzymatic degradation. Each tissue sample was homogenized in 400 µL of ice-cold methanol to extract neurotransmitters. To ensure analytical consistency, isoprenaline was added as an internal standard, and the mixture was vortexed thoroughly to ensure uniform blending. Following homogenization, samples were centrifuged at 12,000 rpm for 10 minutes at 4°C to separate the supernatant from the pellet. The supernatant, containing soluble metabolites, was collected, and the pellet was vacuum-dried for 2 hours to remove residual moisture. The dried pellet was then reconstituted in 200 µL of methanol, vortexed to ensure a uniform solution, and centrifuged again at 12,000 rpm for 5 minutes. The resulting clear supernatant was transferred into autosampler vials for further analysis. Quantification of neurotransmitters, including noradrenaline and serotonin, was carried out using liquid chromatography-mass spectrometry (LC-MS). Calibration curves for each analyte were prepared using standard solutions at concentrations of 100, 50, 25, 12.5, 6.25, and 3.125 ng/mL to enable accurate quantification across a broad dynamic range. Separation was performed on a Thermo Hypersil GOLD™ C18 column, which provided high-resolution chromatographic separation. The mobile phase consisted of 0.1% formic acid in water (solvent A) and acetonitrile (solvent B), using a gradient elution strategy to optimize analyte separation. A 20 µL injection volume and a flow rate of 0.3 mL/min were used throughout the analysis. Detection was performed using targeted selective ion monitoring (tSIM) with a heated electrospray ionization (HESI) source, which enabled sensitive and selective detection of the target compounds. The HESI source was optimized with a vaporizer temperature of 320°C and an ion transfer tube temperature of 300°C. Sheath, auxiliary, and sweep gases were set to 40, 15, and 1 arbitrary unit, respectively, to maintain spray stability and improve ionization efficiency. The spray voltage was maintained at +3500 V in positive ion mode. Additional instrument settings, including resolution, microscans, scan range (m/z), and RF lens voltage, were optimized to enhance the accuracy and sensitivity of tSIM acquisition. Data acquisition and processing were conducted using Xcalibur Qual Browser version 4.4 (Thermo Fisher Scientific), which enabled the reliable identification and quantification of neurotransmitters. The optimized calibration, instrument conditions, and analytical workflow provided robust quantification of glutamine, glutamate, and GABA in the mPFC, contributing valuable insights into regional neurotransmitter regulation (He et al., 2013; Kesharwani et al., 2025a; Kesharwani et al., 2025b; Lim et al., 2023).

**Results:**

**Effect of aging on total exploration time in TO, OiP and NOR:** One-way ANOVA was performed to assess the impact of aging on total exploration time across three behavioral paradigms: TO, OiP, and NOR. The results revealed no statistically significant differences in total exploration time among age groups in any of the tasks (TO: F_2, 27_ = 0.1552, p = 0.8570; OiP: F_2, 27_ = 3.196, p = 0.0568 and NOR: F_2, 27_ = 2.120, p = 0.1395). These findings suggest that aging did not significantly alter the animals’ general exploratory behavior or locomotor activity during task performance.

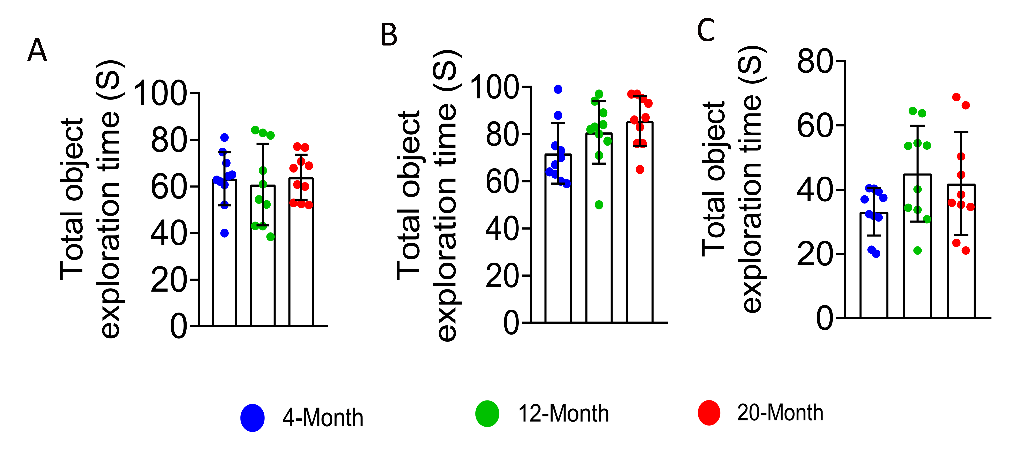

**Fig. S2:** Total exploration time during the behavioral tasks revealed no significant differences between groups, indicating that treatment or aging did not influence overall exploratory activity in TO ***(A)***, OiP ***(B)*** and NOR ***(C)***. Data are presented as mean ± SEM and analyzed using one way ANOVA following Tukey’s multiple comparison test. N = 10 mice per group.

**Effect of CBD on total exploration time across aging groups in TO, NOR, and OiP tasks:** To evaluate the effect of cannabidiol (CBD) treatment on total exploration time across aging groups, unpaired t-tests were conducted for each behavioral task TO, OiP, and NOR. The analysis revealed no statistically significant differences in total exploration time between CBD-treated and aged animals in any of the tasks (TO: p = 0.7651; OiP: p = 0.6451; NOR: p = 0.4498). These results indicate that CBD administration did not influence general exploratory behavior or locomotor activity.

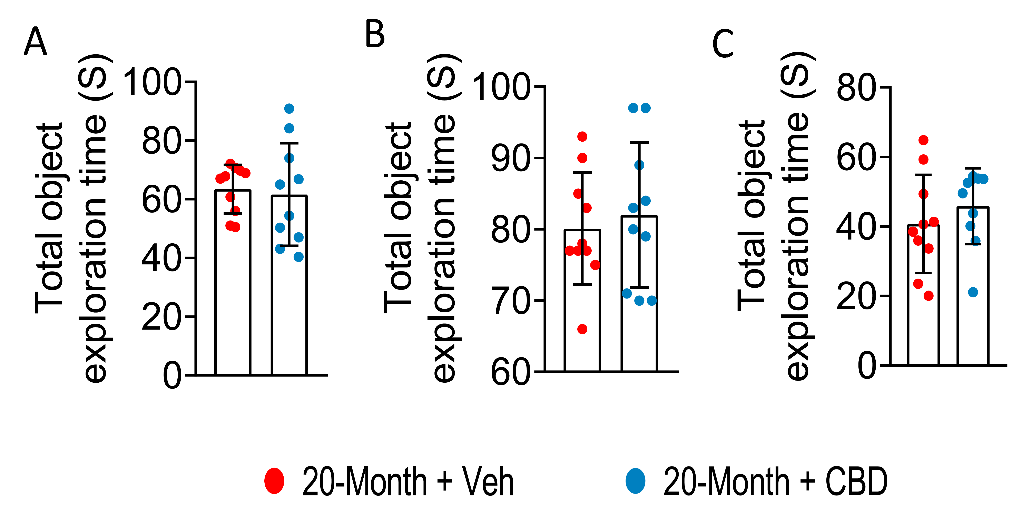

**Fig. S3:** Total exploration time measured during the TO ***(A)***, OiP ***(B)***, and NOR ***(C)*** tasks revealed no significant differences between groups. These results indicate that neither aging nor CBD treatment affected overall exploratory activity. Data are presented as mean ± SEM and analyzed using unpaired t-test. N = 10 mice per group.
